# Molecular basis of AMPA receptor labeling by ligand-directed acyl imidazole chemistry in living neurons

**DOI:** 10.64898/2026.08.31.748281

**Authors:** Dulce C. Guzmán-Ocampo, David De Sancho, Xabier López

## Abstract

Rational design of covalent protein-labeling reagents in complex biological environments requires a molecular-level understanding of how the protein microenvironment governs chemical reactivity; yet, such mechanistic details remain inaccessible to experimental methods alone. In living neurons, Ligand-Directed Acyl Imidazole (LDAI) chemistry has been used to label AMPA receptors as a traceless, affinity-based protein labeling method. Although LDAI labeling reagents have been optimized in the lab, the atomic details of their interactions with the protein and the underlying mechanism remain elusive. In this work, we combined Quantum Mechanical (QM) calculations and molecular dynamics (MD) simulations to propose a detailed reaction mechanism for AMPAR labeling by LDAI reagents and to clarify how the protein microenvironment governs reactivity. Although Lys residues are usually protonated at physiological pH and therefore less nucleophilic in water, our QM results show that Lys labeling is energetically more favorable than competing reactions with Ser or water. MD simulations reveal that PFQX —the LDAI reagent precursor— binds dynamically to the GluA2 AMPAR as an antagonist, inducing conformational changes that reshape the local environment of the acyl imidazole (AI) warhead, underscoring that ligand identity strongly affects labeling outcomes. We also identified intra and intermolecular hydrogen bond networks that may contribute to further immobilize and pre-organize the LDAI reagent. Moreover, the probe’s chemical nature shapes its interactions with the Ligand Binding Domain (LBD), offering a plausible rationale for the previously experimentally observed ligand-dependent fluorescent response. Taken together, our results establish design principles for exploiting the reagent geometry and binding pocket hydrogen-bonding networks for the rational design of LDAI reagents.

## Introduction

The AMPA-type glutamate receptor (AMPAR) is the primary mediator of fast excitatory neurotransmission in the vertebrate central nervous system.^1–4^ These receptors are fundamental to synaptic plasticity, the cellular process underlying learning, memory, and cognition.^2,5–9^ Accordingly, AMPAR dysregulation is strongly linked to numerous neurological, neurodegenerative, and psychiatric disorders, including Alzheimer’s disease, epilepsy, depression, and schizophrenia.^2,4,10,11^ Structurally, AMPARs are organized as heterotetramers, assembled from four distinct subunits: GluA1, GluA2, GluA3, and GluA4 (see Figure 1A).^4,11–13^ While subunit composition varies by brain region —for instance, GluA1/A2 and GluA2/A3 predominate in hippocampal CA1 neurons—, GluA2 is the most abundant subunit across the brain^3,9,14,15^ and has been the primary target of recent affinity-based labeling approaches.^2–4,9,16,17^

**Figure 1:**
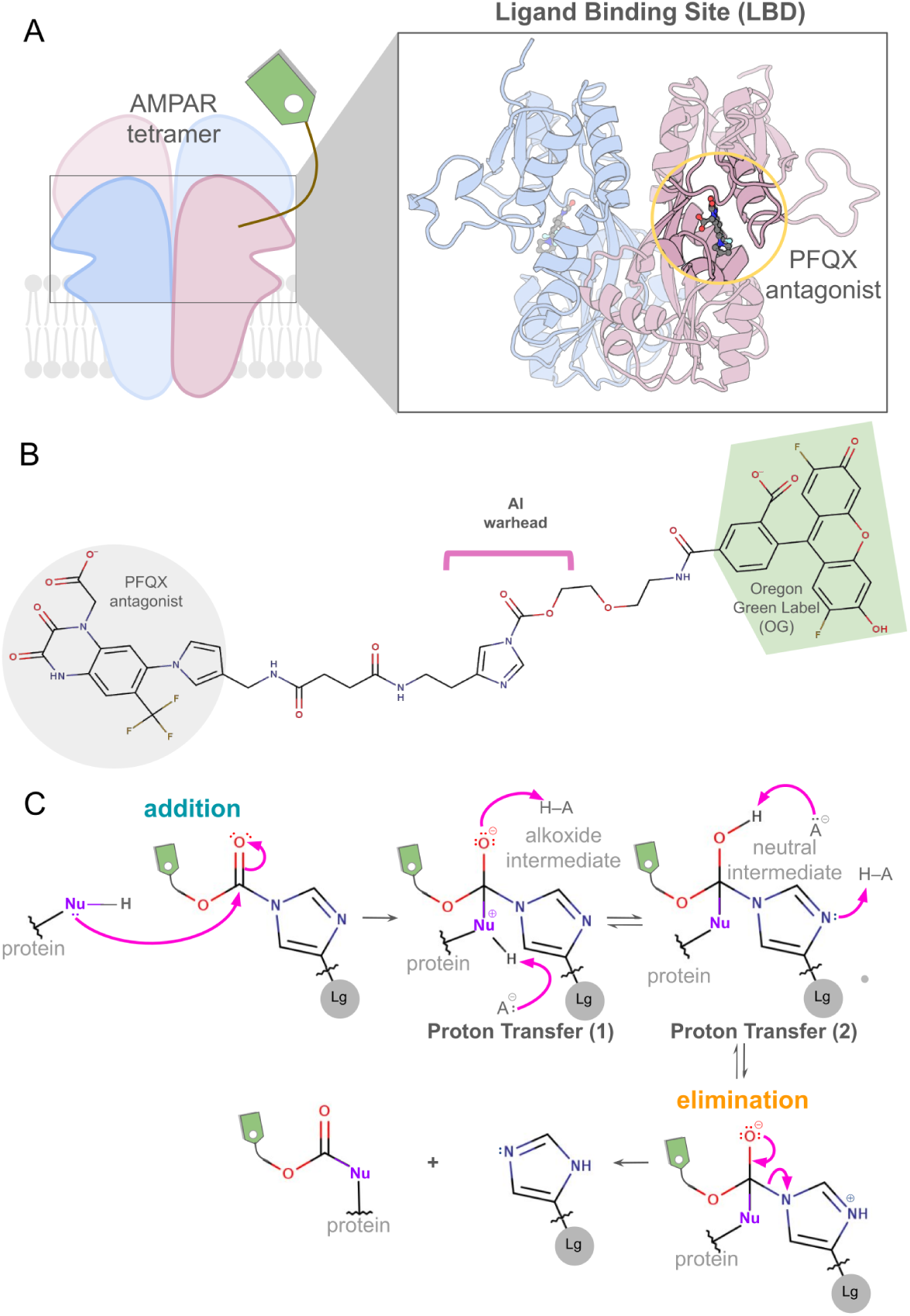
A) Cartoon representation of the AMPAR and PFQX antagonist bound to the ligand binding domain (LBD). B) Depiction of CAM2(OG) labeling reagent. C) Proposed reaction mechanism for the acyl transfer mechanism that lead to the AMPAR labeling.

Ligand-Directed Acyl Imidazole (LDAI) chemistry is an affinity-driven protein labeling method designed to selectively modify the surface of a protein of interest (POI) without disrupting its native biological function.^8,18^ This strategy facilitates the study of endogenous brain receptor dynamics in their native cellular environment without the need for genetic manipulation of the target.^4,11^ LDAI has been used to map the 3D distribution of endogenous neurotransmitter receptors, such as AMPAR, NMDAR, mGlu1, and GABA*_A_*R, in healthy and diseased mouse brains.^11^ Additionally, this chemistry was used to tether fluorescein to folate receptors (FRs) in live KB cells. The labeled receptors acted as “turn-on” biosensors that retained their ability to bind natural ligands.^8,19^

Covalent protein-targeting strategies fall into two broad classes: those designed to label a protein without perturbing function, such as LDAI, and those intended to durably perturb biological activity.^20,21^ The field has evolved from an early focus on highly nucleophilic cysteines towards a vast chemical toolkit capable of modifying nearly every amino acid side chain.^20,22–24^ Selectivity is achieved by matching the intrinsic nucleophilicity or reactivity of a residue with a specific electrophilic “warhead”, ^20,25,26^ often modulated by the protein’s local microenvironment, which can drastically alter the residue’s p*K*_a_ and accessibility.^20,27–29^ A central challenge in directed covalent targeting is finding the balance between a “warhead” that is reactive enough to form a bond but selective enough to avoid off-target toxicity.^20,24–26^ This balance is critical: while high reactivity ensures efficient target engagement, it often leads to promiscuous reactions with abundant off-target nucleophiles, whereas excessive selectivity might result in insufficient potency or slow reaction kinetics.^20,22,24^ LDAI chemistry navigates this challenge through proximity-driven reactivity, exploiting ligand affinity to concentrate a mildly electrophilic warhead at the target site.

The LDAI process proceeds in three steps. First, a high-affinity ligand for the target protein, connected to a functional probe via an acyl imidazole linker, binds reversibly to the protein’s active or binding site. ^8,18^ In the case of AMPAR, the PFQX antagonist that binds to its Ligand Binding Domain (LBD) is used as a seed to build the CAM2(OG) labeling reagent (see Figures 1A and B). Once bound, the acyl imidazole moiety, a moderately reactive electrophile, is positioned in proximity to nucleophilic amino acid residues on the protein’s surface, facilitating an acyl transfer reaction.^2,8,19^ Finally, after the probe is covalently tethered, the ligand-imidazole conjugate is released from the binding pocket, allowing the protein to retain its original activity and be studied in its natural physiological state.^8,18^ Figure 1C illustrates the addition-elimination reaction mechanism that the protein nucleophile (Nu) and LDAI reagent undergo. First, the nucleophile adds to the acyl group, forming a locally zwitterionic tetrahedral intermediate. Subsequent proton transfer (PT) steps, modulated by pH and neighboring acid-base residues, protonate the imidazole ring. This results in the elimination of the ligand-imidazole conjugate and the formation of a carbamate or carbonate modified nucleophile, depending on whether the nucleophilic atom covalently attached to the probe is nitrogen or oxygen.^19,30,31,31–33^ This acyl transfer reaction follows pseudo-second-order kinetics, with rate constants (10^1^ to 10^2^ M^-1^s^-1^) comparable to those of the copper-catalyzed azide-alkyne cycloaddition (CuAAC) click reaction.^8,18^ Correct orientation of the AI warhead to a target nucleophile, enhanced by specific interactions like hydrogen bonding, further contributes to the observed rate.^18,34^

Although the LDAI reaction primarily targets Lysine (Lys) residues, it can also react with Serine (Ser) and Tyrosine (Tyr). ^8,18^ For example, in GluA2 the primary labeling site has been identified as Lys470.^4^ The length and flexibility of the linker between the ligand and the reactive site are critical design parameters, as they determine whether the AI warhead can reach a target nucleophile on the protein surface.^4,11^ There has been much progress recently in LDAI reagent design, including its extension to anchor “clickable” handles such as trans-cyclooctene, (TCO) onto a receptor for subsequent tetrazine ligation.^3,17^ However, the molecular determinants of successful labeling remain poorly understood. It is unclear which surface-exposed nucleophiles are competent labeling targets, why Lys is preferentially modified despite being predominantly protonated at physiological pH, and whether neighboring residues are required to interact with the LDAI reagent to promote the acyl transfer reaction. Addressing these questions is essential for the rational design of reagents with improved selectivity and broader applicability; yet, the atomic-level mechanistic detail required is beyond the reach of experiments alone. In this study, we integrated DFT calculations with classical MD simulations to characterize, at atomic resolution, the reactivity involved in the AMPAR labeling mechanism, along with the interactions that govern it and, consequently, the protein microenvironment that contributes to the reaction. Specifically, we computed reaction energy profiles to confirm the preferred nucleophilic target and the role of bioconjugation-site residues in modulating labeling barriers, characterized the dynamic binding mode of the seed-antagonist PFQX to GluA2 AMPAR and identified the hydrogen-bonding interactions responsible for LDAI reagent immobilization and pre-organization. Together, these results provide a mechanistic framework for the structure-guided optimization of LDAI reagents.

## Results and discussion

### QM Characterization of the LDAI Labeling Mechanism

To establish a baseline mechanistic picture, we first examined the intrinsic reactivity of the LDAI warhead toward different nucleophiles using truncated QM model systems in the absence of the full protein environment. In the case of CAM2(OG), where the “OG” stands for the Oregon Green fluorescent probe, labelling of AMPAR was distributed at 74% on Lys470(Lys60), 11% on Ser424(Ser14), and 15% on Ser673(Ser140).^4^ Throughout this work, we refer to the CAM2(OG) reagent simply as CAM2 and we use the AMPAR LBD numbering as shown in Table S1.

We first investigated the Lys labeling mechanism using truncated model systems representing Lys and CAM2. To this end, we performed QM calculations at the CAM-B3LYP/def2SVP^35–37^ level of theory with D3 dispersion correction^38,39^ and water solvation via the polarizable continuum model (PCM),^40^ supplemented by explicit water molecules as needed to mediate the proton transfer steps, following the reaction scheme in Figure 1C. The resulting energy profile of the reaction is shown in Figure 2. First, the amine nucleophile adds to the acyl group, forming a tetrahedral intermediate (Int^+/-^) in which the amine group and alkoxide oxygen carry formal charges. Int^+/^undergoes a water-mediated 1,3 proton transfer to yield the hemiaminal Int^0^. Elimination of the imidazole ring (Im) from Int^0^ yields the carbamate product and Im-antagonist conjugate (Prod). The carbamate’s protonated acyl group at Prod exergonically transfers its proton to the eliminated Im ring, yielding the fully deprotonated carbamate and the protonated Im-antagonist conjugate in Prod_2_.

**Figure 2:**
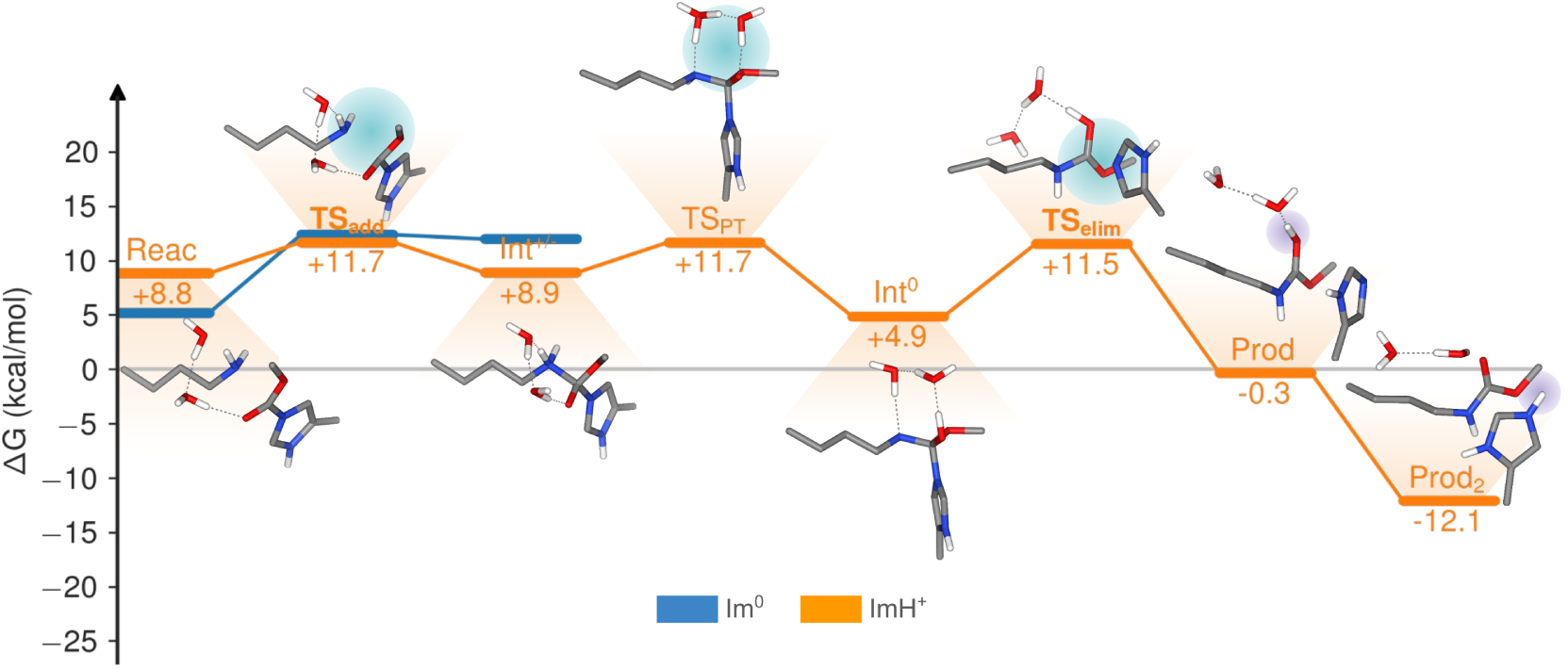
Energy profile of Lys labeling reaction mechanism. The cyan shades highlight the breaking and formation of bonds, and the violet shades, the transfer of a single proton. For clarity, only the addition step for the Im^0^ ring state is shown here, as the ImH^+^ state exhibits a lower energy profile; the full profile is provided in Figure S1.

At physiological pH of 7.4, approximately 99.9% of amino groups in Lys residues are protonated, which normally results in very low nucleophilic reactivity.^18,34^ Despite this, the LDAI reaction remains efficient due to the proximity effect that allows the acylation reaction to proceed even under the unfavorable protonation conditions.^8,18^ Activating Lys nucleophilicity requires accounting for a deprotonation penalty, calculated according to

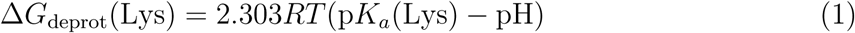

The protonation state of the imidazole ring (Im^0^ or ImH^+^) must also be considered, as it influences both the reaction pathway and the associated energy barriers. The Δ*G*_prot_ in Tables 1, 2, 3, S2 and S4 represents the sum of Δ*G*_deprot_(Lys) of Lys and Δ*G*_prot_(Im) of the imidazole ring, depending on the protonation states considered. Since the p*K_a_* for the Im ring in CAM2 or its cropped system is not reported, we estimated a p*K_a_* value of 4.33*±*0.14 by performing the linear regression presented in Figure S2, detailed in the Methods section. To build the energy profile, we additionally evaluated the number of explicit water molecules required and assessed whether elimination could occur directly after addition, bypassing the proton-transfer step, following an addition*→*elimination rather than addition*→*PT*→*elimination pathway. The resulting barriers for all combinations of these variables are presented in Table S2.

**Table 1:** Comparison of energy barriers of transition states (TS) for Lys and Ser labeling and CAM2 hydrolysis. The numbers (1), (2), and (3) indicate the sequential order of the mechanistic steps. With bold style is highlighted the higher energy barrier and gray fill indicate the lowest energy path between both elimination pathways. ΔG_prot_ values are added to the TS barriers.

| System | $\Delta G_{\text{prot}}$ | TS<sub>add</sub> (1) / TS<sub>add-PT</sub> (1) | TS<sub>elim</sub> (2) | |
| --- | --- | --- | --- | --- |
|  |  |  | TS<sub>PT</sub>(2) | TS<sub>elim</sub>(3) |
| Lys·CAM2(ImH<sup>+</sup>)·2H<sub>2</sub>O | +8.8 | **11.7** | 12.8 |  |
|  |  |  | 11.7 | 11.5 |
| Ser·CAM2(ImH<sup>+</sup>)·2H<sub>2</sub>O | +3.6 | **17.5** | 12.4 |  |
| CAM2(ImH<sup>+</sup>)·3H<sub>2</sub>O | +3.6 | **17.5** | 12.4 |  |

**Table 2:**
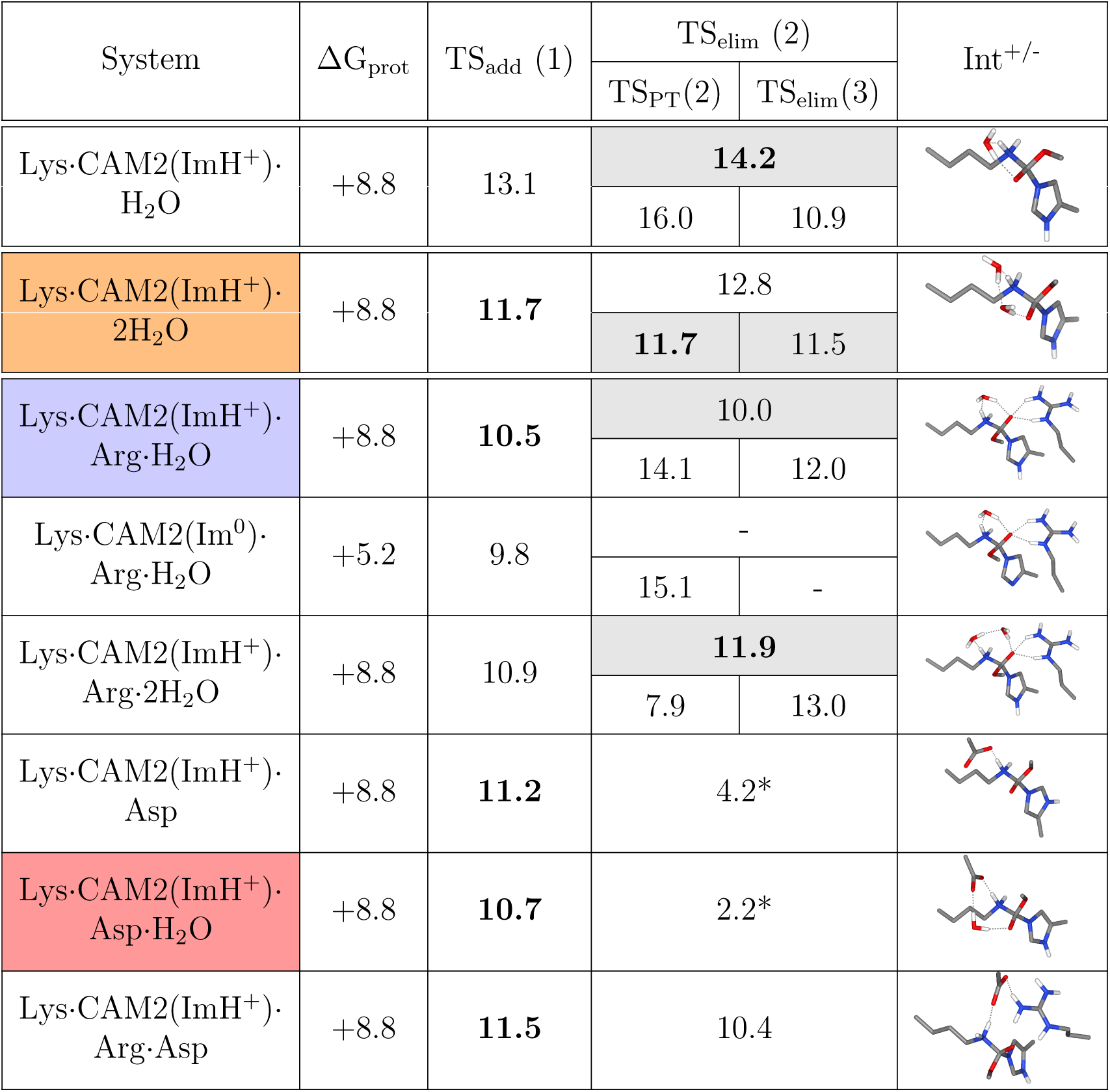
Energy barriers of transition states for Lys labeling assisted by Arg and Asp. References are each respective reactive complex. Units are in kcal/mol. The gray fill highlights the lowest energy path, if applicable, and the bold formatting indicates the highest energy barrier. The numbers (1), (2), and (3) indicate the sequential order of the mechanistic steps. * denotes the energy barrier associated with transfer of the Lys proton to the Asp that leads to the elimination.

| System | $\Delta G_{\text{prot}}$ | TS<sub>add</sub> (1) | TS<sub>elim</sub> (2) | | Int<sup>+/-</sup> |
| --- | --- | --- | --- | --- | --- |
|  |  |  | TS<sub>PT</sub>(2) | TS<sub>elim</sub>(3) |  |
| Lys·CAM2(ImH<sup>+</sup>)·H<sub>2</sub>O | +8.8 | 13.1 | **14.2** |  |  |
|  |  |  | 16.0 | 10.9 |  |
| Lys·CAM2(ImH<sup>+</sup>)·2H<sub>2</sub>O | +8.8 | **11.7** | 12.8 |  |  |
|  |  |  | **11.7** | 11.5 |  |
| Lys·CAM2(ImH<sup>+</sup>)·Arg·H<sub>2</sub>O | +8.8 | **10.5** | 10.0 |  |  |
|  |  |  | 14.1 | 12.0 |  |
| Lys·CAM2(Im<sup>0</sup>)·Arg·H<sub>2</sub>O | +5.2 | 9.8 | - |  |  |
|  |  |  | 15.1 | - |  |
| Lys·CAM2(ImH<sup>+</sup>)·Arg·2H<sub>2</sub>O | +8.8 | 10.9 | **11.9** |  |  |
|  |  |  | 7.9 | 13.0 |  |
| Lys·CAM2(ImH<sup>+</sup>)·Asp | +8.8 | **11.2** | 4.2* |  |  |
| Lys·CAM2(ImH<sup>+</sup>)·Asp·H<sub>2</sub>O | +8.8 | **10.7** | 2.2* |  |  |
| Lys·CAM2(ImH<sup>+</sup>)·Arg·Asp | +8.8 | **11.5** | 10.4 |  |  |

**Table 3:**
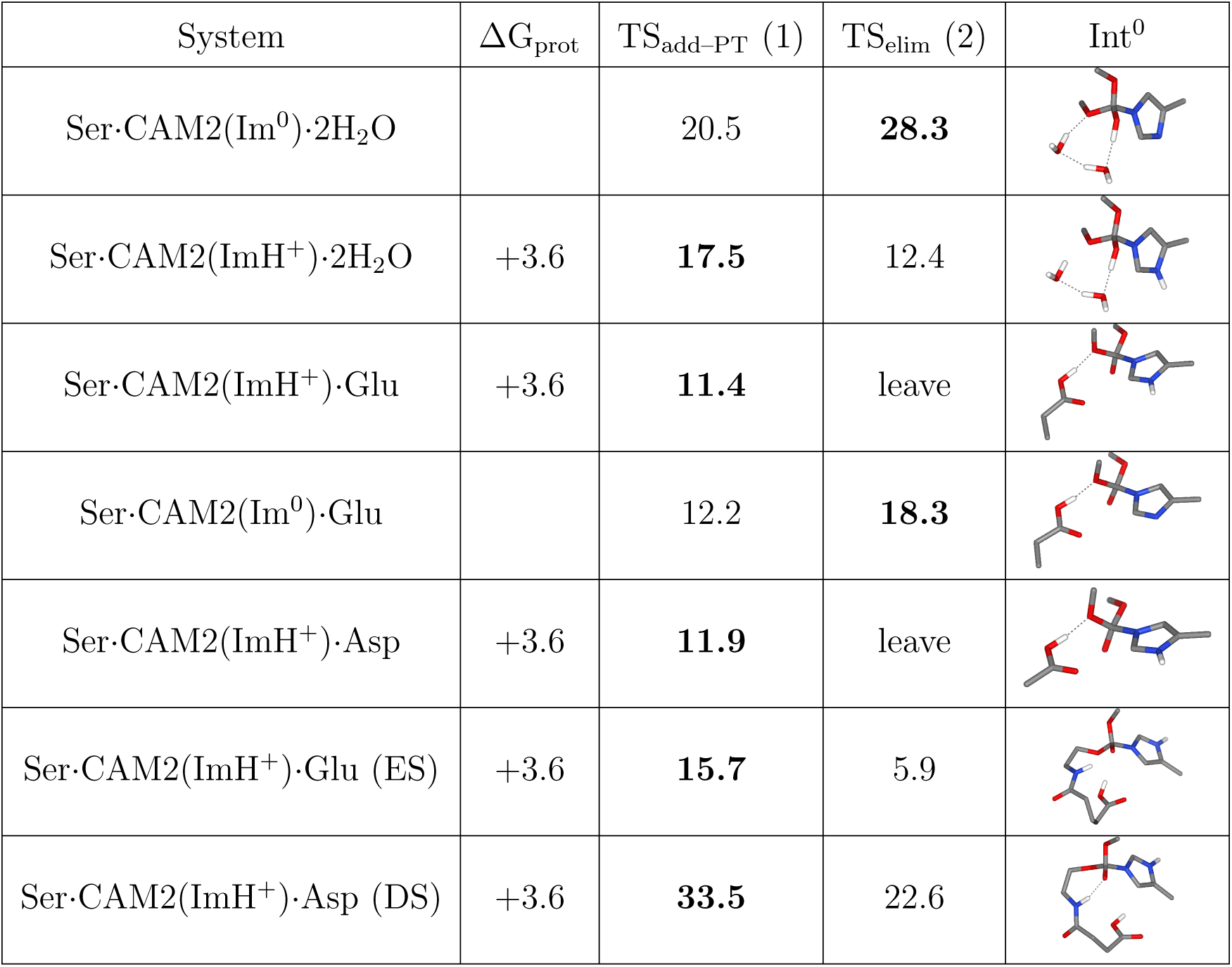
Energy barriers of transition states for Ser labeling assisted by Glu and Asp. References are each respective reactive complex. Units are in kcal/mol. Bold formatting indicates the highest energy barriers. The numbers (1), (2), and (3) indicate the sequential order of the mechanistic steps.

| System | $\Delta G_{\text{prot}}$ | TS<sub>add-PT</sub> (1) | TS<sub>elim</sub> (2) | Int<sup>0</sup> |
| --- | --- | --- | --- | --- |
| Ser·CAM2(Im<sup>0</sup>)·2H<sub>2</sub>O |  | 20.5 | **28.3** |  |
| Ser·CAM2(ImH<sup>+</sup>)·2H<sub>2</sub>O | +3.6 | **17.5** | 12.4 |  |
| Ser·CAM2(ImH<sup>+</sup>)·Glu | +3.6 | **11.4** | leave |  |
| Ser·CAM2(Im<sup>0</sup>)·Glu |  | 12.2 | **18.3** |  |
| Ser·CAM2(ImH<sup>+</sup>)·Asp | +3.6 | **11.9** | leave |  |
| Ser·CAM2(ImH<sup>+</sup>)·Glu (ES) | +3.6 | **15.7** | 5.9 |  |
| Ser·CAM2(ImH<sup>+</sup>)·Asp (DS) | +3.6 | **33.5** | 22.6 |  |

This reaction energy profile for Lys labeling, shown in Figure 2, features three transition states associated with the addition (TS_add_), proton-transfer (TS_PT_), and elimination (TS_elim_) steps. The highest barrier value is 11.7 kcal/mol, common to TS_add_ and TS_PT_, both associated with the formation of the hemiaminal intermediate Int^0^, identifying these two steps as jointly rate-determining. The remaining barriers are lower, indicating that once the hemiaminal is formed, elimination proceeds downhill. These results establish the intrinsic energetic cost of Lys acylation by the AI warhead in the absence of protein microenvironment effects, providing the baseline against which residue-assisted mechanisms will be assessed in subsequent sections.

Systematic variation of the number of explicit water molecules in the Lys*·*CAM2(ImH^+^) model reveals that two water molecules are sufficient to minimize TS_add_, with additional waters providing no further reduction in barrier height. With two explicit water molecules, Im protonation is found, as expected, to enhance reactivity, outweighting the cost of Im protonation by reducing the TS_add_ barrier by 0.7 kcal/mol. Even though, as shown in Figure 2, this is a slight difference, the relative Δ*G* values indicate that the Im ring in Int^+/^is more likely to be protonated, in contrast to the reactive complexes. This means that even if the addition happened with Im^0^, the Int^+/-^(Im^0^) would protonate to Int^+/-^(ImH^+^). Taken together, the TS barriers in Table 1 allow us to identify that Lys*·*CAM2(ImH^+^)*·*H_2_O is more likely to follow the addition*→*PT*→*elimination path than the addition*→*elimination.

### Intrinsic Nucleophile Selectivity in Aqueous Solution

When we consider Ser or water instead of Lys as the nucleophile, the change from nitrogen to oxygen renders the addition and proton-transfer steps concerted, here denoted as an addition—PT step. Consistent with previous reports on acyl imidazole hydrolysis, ^32,41^ three water molecules were included in the hydrolysis model, with one water molecule serving as the nucleophile in place of Lys. As observed for Lys, Im protonation reduces the addition—PT barrier sufficiently to compensate for the protonation cost (Figure S3). The calculated addition—PT barrier for the hydrolysis is 17.5 kcal/mol, i.e. 5.8 kcal/mol higher than the Lys reaction. This is consistent with experimental data indicating that labeling strongly competes with hydrolysis: although the LDAI warhead is relatively stable toward hydrolysis (hydrolysis half-life of 24 hours at pH 6.0), ^8,19^ it undergoes substantially faster labeling in the living brain (labeling half-life of 2–4 hours). ^11^ Table 1 compiles the optimal energy barriers for Lys and Ser labeling in aqueous solution, as well as for CAM2 hydrolysis. The highest energy barrier associated with Ser labeling is 17.5 kcal/mol, the same as that for CAM2 hydrolysis. Since these values exceed the barrier for the Lys labeling reaction by a factor of about 1.5, they rationalize the experimentally observed preference for Lys modification. However, Ser labeling does occur experimentally, accounting for a total of 31% of labeling events. This finding suggests that the local protein environment contributes to the enhanced labeling competence of Lys60, Ser14, and Ser140, a possibility that we explore using molecular dynamics (MD) simulations (see below).

We additionally carried out single-point energy calculations at the CAM-B3LYP/def2TZVPD level and recomputed the energy barriers for Lys and Ser labeling, as well as for CAM2 hydrolysis. The corresponding data are summarized in Table S3, where we note some discrepancies; notably, in this case, the lowest-energy pathway for Lys*·*CAM2(ImH^+^)*·*H_2_ is the addition*→*elimination, and there is now a gap between the Ser labeling TS_add_ energy barrier and that of CAM2 hydrolysis. Nonetheless, this result remains consistent with the CAM-B3LYP/def2-SVP-only calculations in that both account for the preference for Lys labeling.

### Ligand Binding Dynamics and Warhead Positioning from MD Simulations

Ligand-directed strategies are inherently proximity driven, as ligand binding acts as a molecular anchor that raises the likelihood that the warhead will encounter a target nucleophile, thereby substantially increasing the reagent’s effective local concentration.^8,18^ While PFQX binds with well-defined affinity to the AMPAR LBD, the remainder of the CAM2 molecule, including the AI warhead and fluorescent tag, is not expected to adopt a fixed docked conformation. Consequently, within the combined LBD-solvent space at the interface, the AI warhead and fluorescent tag can explore a spectrum of conformations, constrained only by the molecular geometry of the linker. On the protein side, AMPARs function is coupled to conformational changes that are triggered when the neurotransmitter binds to the LBD.^42,43^ These changes drive domain closure that propagates to the transmembrane region and gates the ion channel.^42,44^ Beyond intrinsic protein dynamics, the conformational state of the LBD is also ligand-dependent, with agonists, antagonists, and partial agonists each stabilizing distinct domain configurations.^43,45^ Understanding how PFQX–LDAI reagent binding reshapes the bioconjugation-site local environment requires explicit simulation of the full reagent–receptor complex under dynamic conditions.

Before characterizing the interaction between GluA2 and CAM2, it is necessary to establish how the binding of this specific antagonist, PFQX, shapes the conformational ensemble of the GluA2—LBD dimer, since this will impact the reach of the flexible CAM2 warhead. To this end, we employed the GluA2(S1S2J) dimer using the sequence reported by Wakayama et al.,^4^ with PDB ID: 3KG2^46^ as the structural template in MODELLER.^47,48^ This template structure contains the AMPAR antagonist ZK 200775, which shares key structural features with PFQX, including the quinoxalinedione scaffold, negatively charged Nsubstituent, and heterocyclic and trifluoromethyl substitutions. The GluA2(S1S2J) dimer has been previously validated as a reliable model of the LBD dimer and is scalable to full-channel events.^49–51^ We performed five replicates of 2 µs of conventional MD simulations of the solvated GluA2(S1S2J)-PFQX dimer at 310 K, matching the physiological temperature, and a constant pressure of 1 bar, using the protocol described in the Methods section.

The evolution of the dimeric conformation and associated functional state was monitored throughout each trajectory to assess the stability of the PFQX binding mode and its effect on LBD dynamics. In Figure 3A, the PFQX bound to a GluA2(S1S2J) monomer is shown, highlighting the residues that are hydrogen bonding with it, and in Figure S4, we show the fractions of these hydrogen bond interactions. We find that the Arg96 side chain consistently interacts with both PFQX quinoxalinedione oxygens, establishing it as the primary anchoring interaction. Using the Arg96–PFQX distance as a proxy for binding stability, PFQX was found to remain stably bound across eight of the ten simulated monomers, with transient displacement of up to *∼*8 °A observed in two cases (Mon B of replicate 01 and Mon A of replicate 05; see Figure S5). In these cases, Arg96 shifted its interaction partner from the PFQX quinoxalinedione oxygens to its N-acetate oxygens, as also shown in Figure S4. In the stable binding mode, the PFQX–GluA2 hydrogen bond network comprises: Arg96(Nη) with both quinoxalinedione oxygens; Thr91 with one quinoxalinedione oxygen; Pro89(N) with the unsubstituted quinoxalinedione nitrogen; and Ser142(N,Oγ) with the N-acetate group (Figure S4). In the two monomers exhibiting binding displacement, the remaining hydrogen bond interactions also weaken concurrently, suggesting that Arg96 anchoring is a prerequisite for the stability of the full PFQX–GluA2 hydrogen bond network.

**Figure 3:**
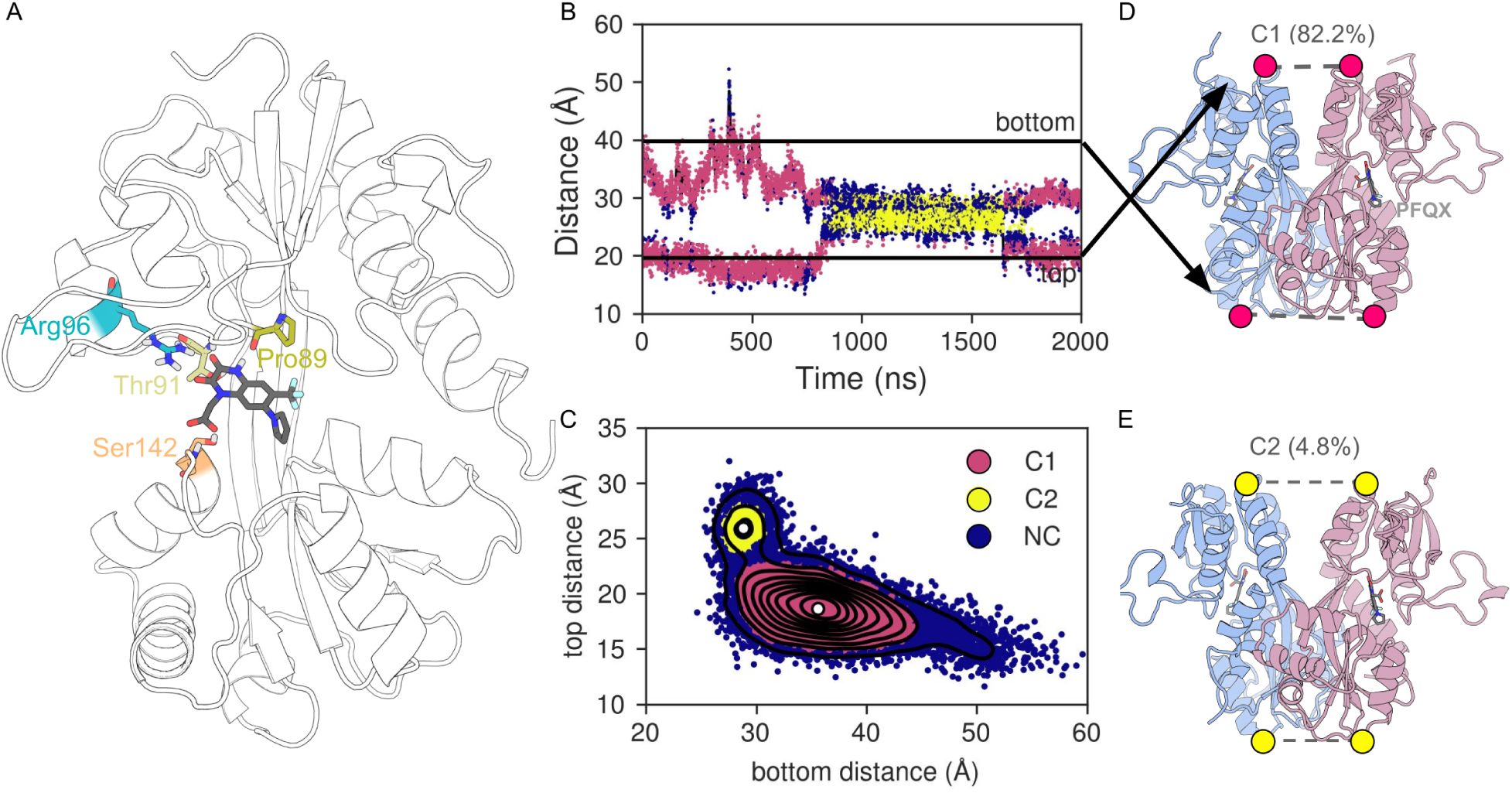
A) Representation of PFQX bound to GluA2(S1S2). B) Detection of the conformational change on GluA2(S1S2)-PFQX dimer in a replicate of our MD simulations. C) 2D clustering to classify all five 2 µs-replicates frames. D,E) Representation of the two conformational changes identified, name as C1 and C2.

It has been previously shown that two LBD-LBD distances — between the monomer-monomer Ser229(Cα) and Lys185(Cα), referred to here as “top” and “bottom” due to their relative positions with respect to the transmembrane region — correlate with the gating mode of the ion channel of the entire AMPAR.^50^ In Figure S6, we show both distances for all five replicates, with initial values indicated as reference. The top distance is more stable across all replicates, while the bottom distance shows greater fluctuations. Comparison with the activated state (free glutamate bound) simulations in Aittoniemi et al., ^51^ reveals comparable top distances of *∼*20 °A. However, bottom distances differ substantially, being 50 °A and above for the activated state, whereas in our GluA2-PFQX simulations, they rarely exceed the initial *∼*40 °A. Following the classification of Hansen et al.,^50^ the initial and predominant conformational state in our simulations corresponds to the resting state. Additionally, a desensitized-like state,^50^ in which the top and bottom distances are more alike, was transiently observed in our replicate 03 (see Figure 3B). Pooling the top and bottom distances across all five replicates and performing a 2D clustering analysis (Figure 3C) revealed two predominant states in our simulations. Representative dimer configurations for each state are shown in Figures 3D and E. We designate these two states as C1 and C2, in order of predominance, with C1 corresponding to the resting-like state and C2 to the desensitized-like state.

To assess the degree of clamshell closure, we monitored the S1-S2 interdomain distance (*d*_S1_*_−_*_S2_) between residues Leu90, Thr91, and Ile92 of the S1 domain and Ser142 and Thr143 of the S2 domain, as defined by Aittoniemi et al. ^51^ In our simulations with the PFQX antagonist, *d*_S1_*_−_*_S2_ is *∼*13.2 °A, compared to *∼*9.3 °A with their free glutamate-bound activated state. Being bulkier than free glutamate, PFQX should not cause clamshell closure and, therefore, channel opening, which is consistent with its acting as an antagonist. As shown in Figure S7, two notable observations emerge: (i) the PFQX binding mode shift causes the clamshell closure to *∼*9.7 °A, approaching the free glutamate bound value, and (ii) the clamshell in one monomer is slightly more open where the C2 desensitized-like state was identified. Both events show a loss of the PFQX-Ser142 hydrogen bond interaction in Figure S4, but in the case of the C2 state, there is no gain in the PFQX N-acetate–Arg96 interaction. The main-chain *RMSD* between the monomers (Figure S8) reveals that the appearance of the C2 state coincides with monomer differentiation. Taken together, these observations indicate that the conformational differences between C1 and C2 states are driven primarily by monomer differentiation and clamshell opening rather than closure.

### Role of Neighboring Residues in Directing Lys60 Labeling

Having characterized the conformational ensemble sampled by the GluA2–PFQX complex, we next sought to understand how the bioconjugation site and labeling conditions are organized, focusing on the residues that form the conjugation site and using experimental observations that place initial constraints on the labeling geometry. A key experimental observation by Wakayama et al. ^4^ is that one of the four AMPAR subunits, namely GluA1, was not labeled by CAM2.^4^ In GluA1, the two Ser residues labeled in GluA2 are replaced by either Asp (S14D) or Ala (S140A). Since Lys60 is conserved in GluA1 yet labeling is absent, this observation initially suggested that at least one of the Ser residues contributes to Lys60 labeling in GluA2. Nevertheless, the combined sequence differences between GluA1 and GluA2 leave open the possibility that factors beyond the Ser substitutions account for the loss of labeling, underscoring the need for direct structural evidence of Ser involvement.

We therefore examined the proximity of Ser14 or Ser140 to Lys60 in our MD simulations to assess whether either residue is geometrically positioned to participate in the Lys labeling reaction. As both Ser14 and Lys60 belong to the S1 domain, their inter-residue distance is constrained by the tertiary structure of the domain: the backbone-to-backbone distance remains essentially fixed, leaving only the side chains free to change conformations and modulate their proximity. In our GluA2-PFQX simulations, the Ser14-Lys60 side chain distance averaged 19.6*±*2.7 °A and had a minimum at 8.0 °A, ruling out the direct participation of Ser14 in the Lys60 labeling reaction. Ser140, in contrast, belongs to the S2 domain, so its separation from Lys60 is not fixed by the tertiary structure and can vary with interdomain motion. On this basis, and supported by the structural analysis presented below, we propose that Ser140 is the residue that participates in Lys60 labeling. In Figure S9, we show the distances between Lys60 and Ser140 side chains in our GluA2-PFQX dimer simulations, allowing us to identify that they are approximately 11*−*13 °A apart, with transient approaches to within water-bridging distance. Since Lys60 must be deprotonated to act as a nucleophile, as established above, changes in its protonation state may additionally modulate its side chain conformation and proximity to Ser140. Analysis of the Aittoniemi et al. ^51^ trajectories with free glutamate bound reveals that clamshell closure brings Ser140 into hydrogen bonding distance of the Gly62 backbone, an interaction observed in up to 40% of the trajectory frames. Since Gly62 is adjacent to Lys60, this finding indicates that clamshell closure can bring Ser140 and Lys60 into closer proximity than observed in our PFQX-bound simulations.

Inspection of the GluA2(S1S2J) monomer sequence reveals that each labeled nucleophile is immediately preceded by an acidic residue: Lys60, Ser14 and Ser140 belong to the GDGK_60_YG, ES_14_ and DS_140_ motifs, respectively. Acidic residues such as Asp and Glu are known to assist nucleophilic activation as a base or mediate proton transfer in addition-elimination reactions,^52–54^ suggesting that these neighboring residues may play a mechanistic role in LDAI labeling at both the molecular dynamics and QM levels. In our GluA2–PFQX simulations, the Lys60–Asp58 and Ser140–Asp139 pairs approach within hydrogen bonding distance of their respective side chains, whereas the Ser14–Glu13 pair does not (Figures S10, S11, S12). Together, these distance analyzes identify these base-nucleophile pairs as likely participants in the labeling mechanism, a proposition we return to after characterizing the binding dynamics of full CAM2.

### CAM2 Binding Dynamics and Electrophile–Nucleophile Proximity

The fact that labeling occurs experimentally confirms that the AI warhead and target nucleophile can get sufficiently close; however, given the conformational flexibility of CAM2, a single rigid binding pose is not expected. Indeed, our GluA2-PFQX simulations already demonstrated that the PFQX fragment itself binds dynamically, sampling multiple orientations. Starting from the most representative structures of the C1 and C2 conformational states of the GluA2(S1S2J)-PFQX dimer, we generated nine diverse GluA2(S1S2J)-CAM2 starting complexes for each state by building CAM2 from the PFQX scaffold and sampling different conformations. Each of the eighteen GluA2(S1S2J)-CAM2 complexes was equilibrated and subjected to 200 ns of equilibrium MD. As observed in the GluA2(S1S2J)-PFQX simulations, the PFQX fragment of CAM2 exhibited binding fluctuations in some monomers but remained bound throughout all trajectories (Figure S13 and S14). The top and bottom interdomain distances (Figure S15 and S16) reveal that the protein dimer conformations evolved during the simulations, as expected given the dynamic nature of the system. The C2 set exhibited the greatest conformational variation, consistent with its origin from the less populated cluster identified in the GluA2-PFQX analysis. Projecting the top and bottom distances onto the clustering space derived from the GluA2–PFQX analysis (Figure S17) shows that the top distances of both sets shift toward the alternate conformation region, while the bottom distances retain their discriminating power between C1 and C2.

To assess nucleophile accessibility in the CAM2-bound simulations, we monitored the distances between the electrophilic acyl carbon and the nucleophilic atoms of Lys60, Ser140 and Ser14, with the latter represented as a color scale in Figure 4A. This representation reveals a geometric constraint: the proximity of the electrophile to Ser14(Oγ) is mutually exclusive with proximity to either Lys60(Nζ) or Ser140(Oγ). Considered along with the 1D distributions in Figure S18, these data reveal clear differences between the C1 and C2 sets. In the C1 set, the electrophile approaches Ser14(Oγ) with a higher probability and a more compact distance range than in C2. Conversely, the C2 conformation favors the proximity of the electrophile to Lys60(Nζ) compared to C1 and also narrows the accessible distance range to Ser140(Oγ). These findings demonstrate that the protein conformation directly governs both the identity of the residue most accessible to the electrophile and the overall probability of a productive warhead-nucleophile encounter. Although the conformational dependence of nucleophile accessibility is most directly observable in computational studies, it carries practical implications for experimental reagent design: replacing the directing ligand fragment will alter the receptor’s conformational ensemble and consequently shift the distribution of labeling outcomes.

**Figure 4:**
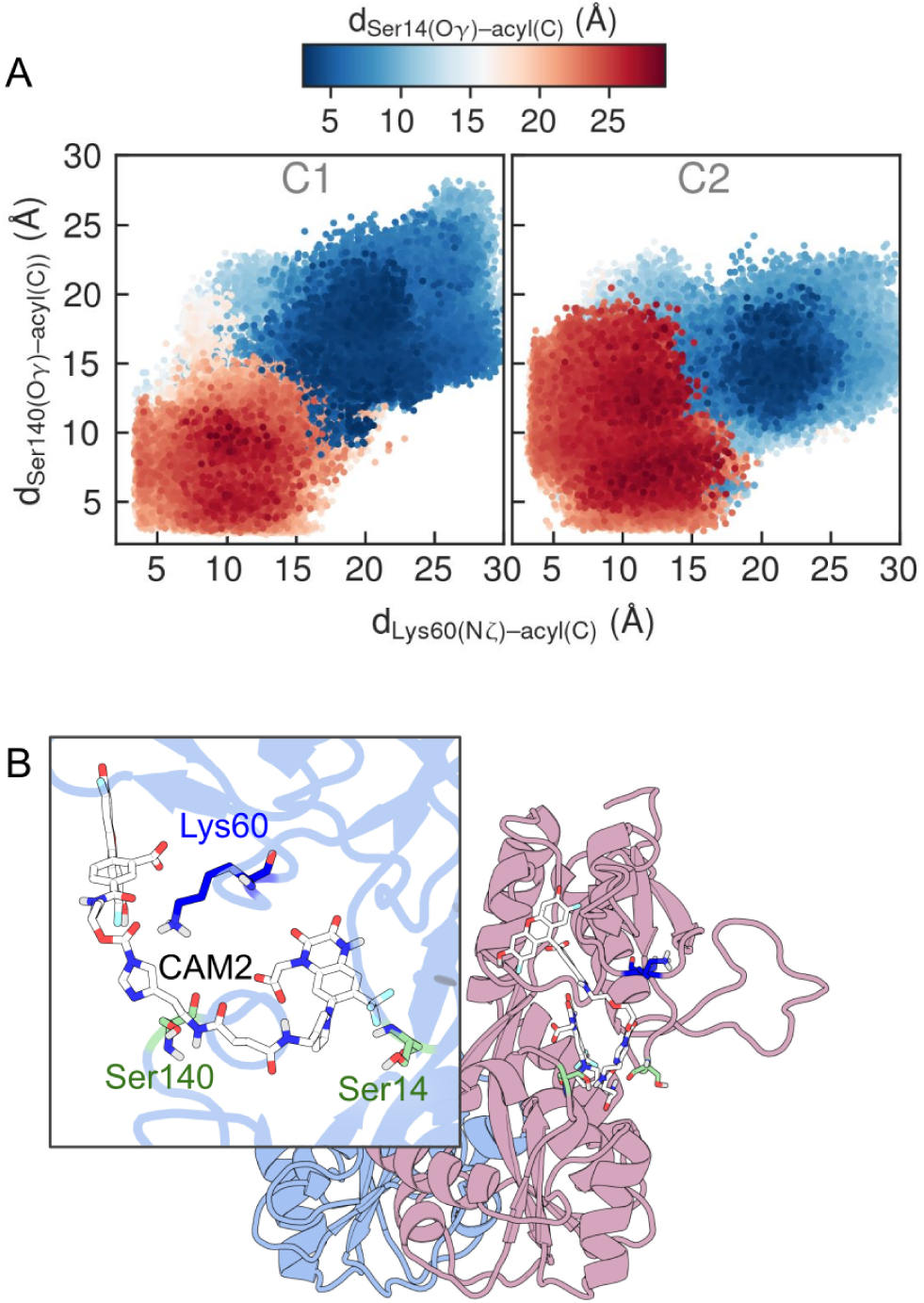
A) 2D distances distributions of CAM2 acyl carbon and Lys60(Nζ) and Ser140(Oγ) of all two sets C1 and C2 and nine 200ns-replicates for the GluA2(S1S2J)-CAM2 dimer system. The respective distances with Ser14(Oγ) is represente by colorscale. B) Representation of CAM2 bound to GluA2(S1S2J), labeled residues Lys60, Ser14 and Ser140 are indicated.

### Protein–CAM2 Interactions and Their Role in Productive Labeling Geometry

To identify the specific protein–reagent interactions that bring the AI warhead into productive proximity with the target nucleophiles, we examined the full GluA2–CAM2 interaction landscape across both conformational sets. Figure S19 extends the analysis of Figure 4A, allowing replicates and monomers to be distinguish, where it is also noted that these distances are not identical in monomer partners. The C2 set exhibits a high density population centered at *d*_Lys60(Nζ)-acyl(C)_,*d*_Ser140(Oγ)-acyl(C)_ = (6.7 °A,9.0 °A), absent in the C1 set, identifying a putative hotspot geometry in which Ser140 forms a hydrogen bond interaction with the second amide group of CAM2 (am2), suggesting that specific protein–reagent hydrogen bonding interactions play a key role in positioning the reactive partners. To characterize these interactions systematically, we analyzed hydrogen bond formation between CAM2 and GluA2(S1S2J) across all eighteen trajectories (two sets × nine replicates), the results of which are presented in Figure S20. The PFQX antagonist fragment of CAM2 maintains hydrogen bond interactions with the LBD consistent with those observed in the GluA2–PFQX simulations. Hydrogen bond interactions between CAM2 and the protein decrease progressively in CAM2’s middle region as it is farther away from the antagonist moiety. The three CAM2 amide groups are designated am1–am3 in order from the antagonist fragment outward. In Figure S20, am1 mostly interacts with Thr174 and Glu13, the residue immediately before labeled Ser14 in the sequence, while am2 interacts with Arg172, Thr173, and Thr174. Hydrogen bond interactions between the GluA2(S1S2J) protein and CAM2 Im, acyl, and am3 are scarce; however, interactions between the labeled nucleophiles Lys60, Ser14, and Ser140 and the Im and acyl groups were nonetheless observed. Notably, Ser140 engages in hydrogen bonding interactions with the am1, am2, Im, and acyl groups of CAM2, which encompass almost the entire central region of CAM2, albeit at low frequency.

The Oregon Green (OG) probe fragment in CAM2, bearing carboxylate, acyl, and hydroxyl groups, as well as an aromatic ring system, engages in hydrogen bonding interactions with the protein. These interactions occur predominantly with Arg and Lys residues, with Arg172 being particularly notable as it simultaneously engages both the OG fragment and the CAM2 middle region. Since the probe region is, by design, modifiable, the specific interactions observed here for CAM2(OG) cannot be considered universal determinants of labeling success across all versions of CAM2 or other LDAI reagents. Nevertheless, since the experimental labeling percentages were obtained with CAM2(OG) specifically, the interactions reported here are directly relevant to this reagent and may serve as a reference point for other similar dyes or probes.

To identify the protein-reagent interactions associated with the productive electrophile-nucleophile proximity, we selected the frames where each labeled nucleophilic atom is within 5 °A of the CAM2 acyl carbon. Residues forming hydrogen bonds with CAM2 in more than 10% of these selected frames are presented in Figure S21. A 2D representation of CAM2 and a reduced version of Figure S21 for Lys60 and the CAM2 central region is presented in Figure 5A,B, showing the interactions that bring the Lys60 amine and CAM2 acyl groups into close proximity. The residues most frequently engaged in hydrogen bond interactions under these conditions are Thr174, Ser140, Arg172, and Lys60 itself, with their specific interactions detailed in Figures 5 and S21. The main difference between the two conformational sets is that Arg172 is the dominant CAM2-interacting residue in C1, while Ser140 takes on this role in C2. In the C1 set, the Thr174*−*X*−*Arg172 tripeptide motif matches with CAM2 am1*−*am2*−*Im groups, forming a complementary linear arrangement of three hydrogen bonds (Figure 5C). At the same time, Arg64 interacts with the OG fragment (Figure S21). Together, these combined interactions immobilize CAM2 in a conformation that brings the Lys60 amine and the CAM2 acyl group into hydrogen bonding distance. In the C2 set, Ser140 replaces Arg172-X-Thr174 as the primary CAM2 partner, with no other residue significantly interacting with CAM2, except for the antagonist moiety. Instead, CAM2 adopts an intramolecular hydrogen bond between its antagonist and OG fragments (Figure 5C), pre-organizing the reagent conformation and facilitating the Lys60(Nζ)-acyl group interaction. This scenario provides a mechanistic basis for Ser140 participation in Lys60 labeling: by simultaneously engaging am1 and am2, Ser140 immobilizes CAM2 and increases the probability of a productive warhead-nucleophile encounter. Both Arg172 and Ser140 form hydrogen bonds with Asp139, which appear to modulate the orientation of the Arg172 guanidinium group towards CAM2 and the availability of Ser140 to interact with CAM2. When Lys60 is positioned near the electrophile, the OG carboxylate group forms hydrogen bonds with Arg148, Arg172, and Arg64 (Figure S21). In the C1 set, interacting with Arg64 orients the OG fragment toward the upper half of the shell, whereas in the C2 set, interaction with Arg148 or Arg172 positions it toward the lower half. Although these interactions are not present in all frames, they suggest that Arg anchoring of the OG-fragment through electrostatic and cation-π interactions contributes to a productive warhead-nucleophile encounter. OG-Arg interactions are also observed when either labeled Ser is positioned near the electrophile (Figure S21); however, when Lys60 is the proximal nucleophile, only Arg residues interact with it directly.

**Figure 5:**
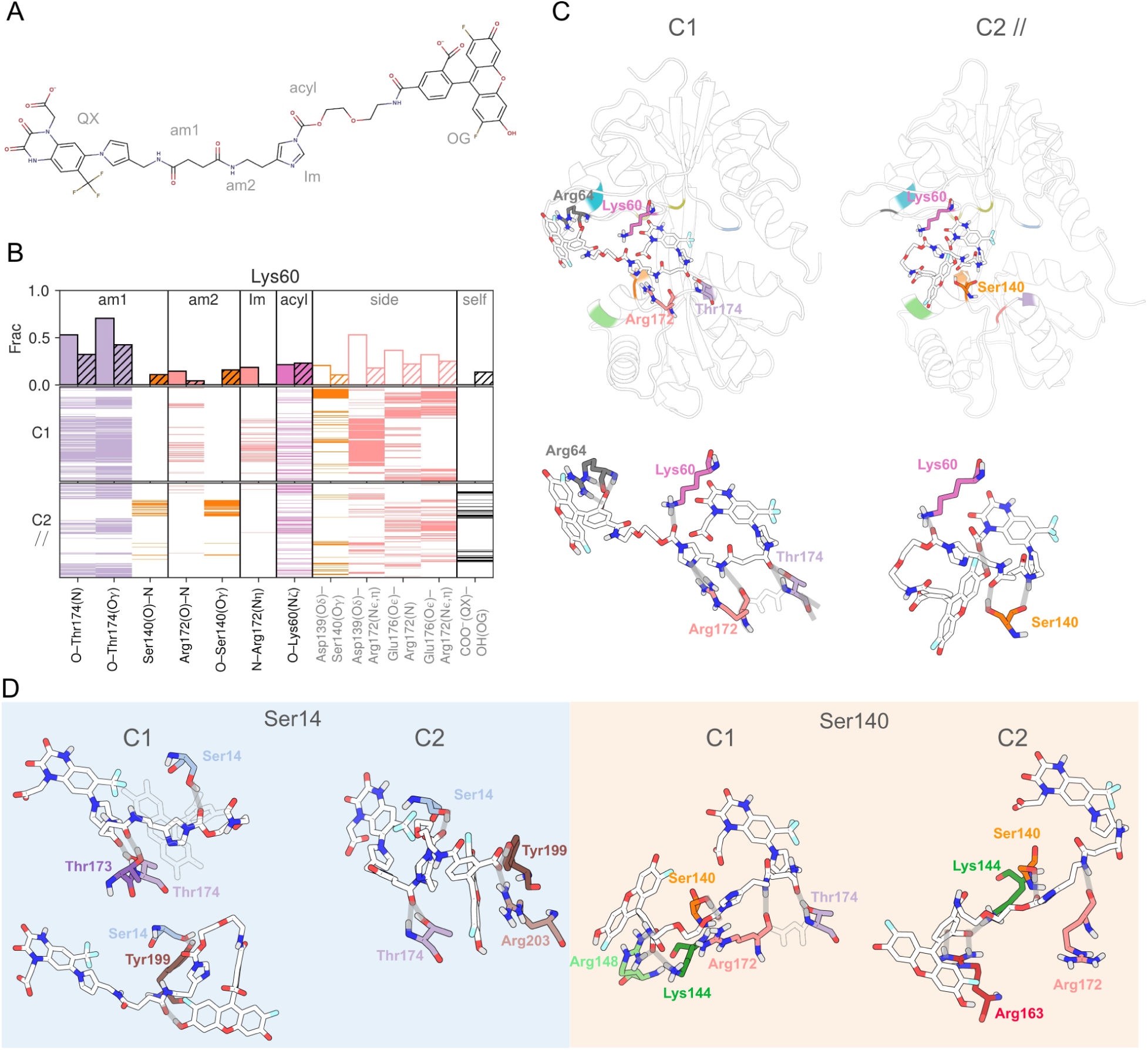
A) 2D representation of CAM2 middle region. B) Barplots of fractions and presences of hydrogen bonds between GluA2(S1S2J) and middle region of CAM2 when Lys60(Nζ) is at least 5 °A from the CAM2 acyl carbon. Interactions are shown from left to right following the CAM2 representation in A). C2 is distinguished from C1 by diagonal hatching (//). Only residues with a fraction *≥* 0.1 are shown. The “side” and “self” sections are filtered side-chain hydrogen bond interactions involving residues that interact with CAM2 and the intramolecular hydrogen bond formed by CAM2, respectively. C) Representations of the CAM2 bound to GluA2(S1S2J) with Lys60 and the AI warhead in close proximity, highlighting the main hydrogen bond interactions observed. D) Selected hydrogen bond interactions between CAM2 and GluA2(S1S2J), with Ser14 and Ser140 indicated as labeling nucleophiles.

When Ser140 acts as the nucleophile, the roles of Arg172 and Asp139 shift in a mechanistically revealing way (Figure 5C,D). In this configuration, the Arg172 backbone and side chain interact only by hydrogen bonding with am2, but not with the Im ring. The guanidiniumIm interaction thus appears to be a key determinant of nucleophile identity, directing the acyl group towards Lys60 rather than Ser140 (Figure 5C). When Ser140 is placed as the nucleophile in Figure S21, Glu13 and Thr174, from different half shells, compete to hold the am1, with Glu13 disrupting the hydrogen bonding with Arg172 and reducing am2 anchoring. The interaction landscape shifts further when Ser14 is positioned as the nucleophile, with Thr173 and Thr174 emerging as the primary am2-interacting residues. The Thr174am1 interaction, the most frequent interaction involving am1 independent of the proximity selection, is common to all three nucleophile cases but occurs more frequently when Lys60 is the nucleophile. Thr173-am2 and Thr174-am2 hydrogen bonding interactions appear to influence the Ser14 placement as a nucleophile, with their occurrence being sensitive to the protein conformational state in the C1 and C2 sets. In the C2 set, when Ser14 is placed as a nucleophile, Arg203 and Tyr199 stabilize the OG-fragment through a combination of electrostatic, cation-π, and π *−* π interactions (Figure 5D). This interaction is enabled by the conformational transition from C1 to C2 observed in GluA2(S1S2J)-PFQX simulations, which disrupts the Arg203-Glu260 salt bridge and frees Arg203 to interact with the OG carboxylate. Notably, Tyr199 —a potential nucleophile— forms hydrogen bonds with Ser14 and is able to reach the acyl group (Figure 5D); yet, it was not identified as a labeled residue by Wakayama et al.,^4^ suggesting that geometric accessibility alone is insufficient to guaranty labeling in the experiments.

Kiyonaka et al. ^2^ reported ligand-induced fluorescent responses in LDAI-labeled receptors, whereby the addition of free glutamate or the NBQX antagonist increases fluorescence intensity.^2^ Our simulations suggest two complementary explanations for this observation: (i) hydrogen bonding between the OG and antagonist fragments, which may persist after covalent probe attachment, and (ii) conformationally induced interactions between the OG fragment and residues such as Arg203 and Glu260. Collectively, these observations underscore that the chemical nature of the probe fragment shapes its interactions with the protein surface in ways that extend beyond the labeling reaction itself, with implications for both reagent design and the interpretation of fluorescence-based readouts.

### QM Characterization of Residue-Assisted Labeling Mechanisms

The GluA2–CAM2 MD analysis identified two distinct interaction motifs, Thr174–X–Arg172 in C1 and Ser140 in C2, that immobilize the CAM2 middle region and promote productive Lys60–electrophile proximity. Furthermore, when Ser140 is not engaged in CAM2 immobilization, Arg172 hydrogen bonding with the Im ring directs the acyl group toward Lys60 rather than Ser140. Asp139 modulates both interactions through hydrogen bonding with Arg172 and Ser140, effectively regulating the balance between the two immobilization motifs. Asp139, the acidic residue immediately preceding Ser140 in the sequence, emerged as a key participant in the hydrogen bond analysis. To assess the participation of the analogous acidic residues preceding Ser14 and Lys60, Glu13 and Asp58, respectively, we quantified their side chain hydrogen bond frequencies in the CAM2-bound simulations. The Asp139–Ser140 pair shows the highest interaction frequency (C1: 8.82%, C2: 25.07%), followed by the water-mediated Glu13–H_2_O–Ser14 interaction (C1: 10.02%, C2: 12.05%), while direct Asp58–Lys60 hydrogen bonding is rare (C1: 0.45%, C2: 5.98%). Although these frequencies are modest, they confirm that all three acidic-nucleophile pairs achieve hydrogen bonding contact, with probabilities that are likely underestimated by classical force fields that do not account for the enhanced reactivity of the bioconjugation site. These observations motivated a return to QM calculations to explicitly test the mechanistic role of the neighboring acid-base residues identified by the MD analysis. Specifically, we examined the effect of Asp139 as an auxiliary base and Arg172 as an oxyanion stabilizer, a role consistent with its known involvement in stabilizing the negative charge on oxyanions,^55,56^ such as the acyl oxygen formed during nucleophilic addition. Table 2 presents the transition state barriers for Argand Asp-assisted Lys labeling.

First we explore the possibility of Arg172 involvement. Arg stabilization of the alkoxide intermediate lowers both TS_add_(1) and TS_elim_(2) in the presence of a single explicit water molecule (Lys*·*CAM2(ImH^+^)*·*Arg*·*H_2_O). This stabilization renders the addition*→*elimination pathway more favorable than the addition*→*PT*→*elimination since the protonation of the alkoxide in the PT step weakens its electrostatic interaction with the positively charged Arg. As a result, the highest energy barrier is 10.9 kcal/mol, 1.2 kcal/mol reduced compared to the unassisted Lys*·*CAM2(ImH^+^)*·*2H_2_O mechanism. Another possibility is the replacement of one water molecule for Asp in the Lys*·*CAM2(ImH^+^)*·*2H_2_O system, which also decreases the energy of TS_add_. Then, Asp acts as a base, taking away the proton from the amine fragment of Int^+/^which leads to a free-barrier elimination taking place. This pathway yields the highest energy barrier of 10.7 kcal/mol, just slightly lower than the Arg-assisted mechanism. The contrasting energy profiles of the optimal mechanism of the reaction of Lys labeling are shown in Figure 6. Finally, a collaborative Arg-Asp mechanism was also explored, yielding a barrier marginally lower than the reference Lys*·*CAM2(ImH^+^)*·*2H_2_O system, but not better than adding either Arg or Asp. This result suggests either that Arg and Asp contribute independently to labeling rather than cooperatively, or that the collaborative mechanism requires more carefully optimized initial geometries than those employed here. Energy barriers listed in Table 2 were also recomputed from single-point energy calculations at the CAM-B3LYP/def2-TZVPD level, and the resulting values are presented in Table S5. The Asp- and Arg-assisted pathways, i.e., Lys*·*CAM2(ImH^+^)*·*Asp*·*H_2_O and Lys*·*CAM2(ImH^+^)*·*Arg*·*H_2_O each consistently lower the highest energy barrier by 1.6 kcal/mol.

**Figure 6:**
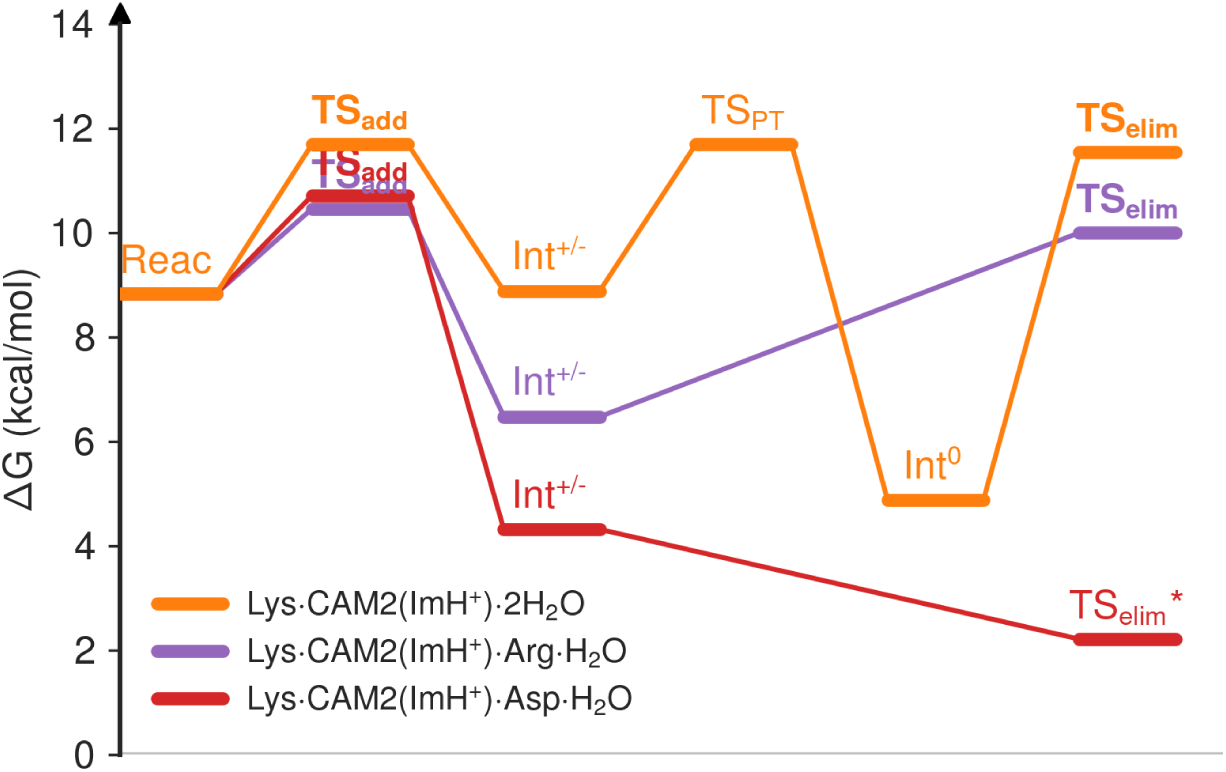
Energy profiles of Argand Asp-assisted mechanisms of Lys labeling reaction. TS_elim_* denotes the transition state where the proton is transferred from Lys to Asp, which triggers the elimination step.

As a completeness check, we assessed whether the protein backbone or Ser residues could favor the reaction by taking on the role of water in stabilizing the addition intermediate and facilitating subsequent elimination. In all cases, the resulting barriers were no lower than those of the reference Lys*·*CAM2(ImH^+^)*·*2H_2_O. The results are reported in Table S4, confirming that neither the backbone nor Ser inclusion provides any energetic advantage over the water-assisted reference. Table 3 presents the energy barriers of Ser labeling assisted by Asp and Glu, modeled using their formate and acetate models, respectively, as residue mimics, or by including the corresponding cropped dipeptides Asp-Ser (DS) and Glu-Ser (ES). Asp/Glu assistance reduces the Ser labeling barrier to a value comparable to unassisted Lys labeling, providing a mechanistic rationale for the experimentally observed Ser labeling. The energy barriers of 11.4 and 11.9 kcal/mol for the labeling of Ser assisted by Glu and Asp, respectively, reflect the well-known fact that Glu is more basic than Asp and thus requires less energy to abstract the proton from the Ser hydroxyl group. Therefore, the energetics contradict the reported Ser labeling percentages: (Glu13-)Ser14 is labeled to a lesser extent than (Asp139-)Ser140, at 11% and 14%, respectively. Thus, this discrepancy may stem from a reduced likelihood of forming an appropriate bioconjugation site configuration for labeling (Glu13-)Ser14 compared with (Asp139-)Ser140. Finally, the increased energy barriers observed when considering the Glu-Ser and Asp-Ser dipeptides, Ser*·*CAM2(ImH^+^)*·*Glu (ES) and Ser*·*CAM2(ImH^+^)*·*Asp (DS), rule out the possibility that the backbone of the acidic residue contributes to facilitating the labeling reaction.

## Conclusions

We combined QM calculations and MD simulations to elucidate the mechanism of AMPAR labeling by LDAI reagents and to establish how the protein microenvironment governs reactivity. QM calculations provided a baseline picture of intrinsic nucleophile selectivity; MD simulations revealed the dynamic binding mode of the PFQX–based LDAI reagent and the hydrogen-bond interactions that immobilize it for productive labeling. Residue-assisted QM mechanisms connected these structural observations to quantitative energy barriers consistent with experimental labeling percentages.

As established by our QM free energy barriers (Table 1), Lys acylation is kinetically preferred over competing reactions with Ser and water, despite being predominantly protonated and, therefore, less nucleophilic in aqueous solution. The intrinsic preference for Lys acylation arises from the addition*→*PT*→*elimination pathway identified as optimal in aqueous solution, with the Im ring protonation state and explicit water molecules playing key roles in modulating the barrier heights. We also found that the PFQX antagonist, and consequently the LDAI reagent derived from it, binds dynamically to the GluA2 AMPAR, inducing conformational transitions in the protein that may alter the microenvironment around the AI warhead. This observation highlights how crucial the identity of the ligand seed is and how modifying it can alter labeling outcomes. Far from just being a juxtaposition, as had been previously noted, hydrogen bond interactions drive labeling efficiency.^34^ Either the Arg172-X-Thr174 motif or Ser140 forms hydrogen bond interactions with the amides or the acyl imidazole of the LDAI reagent to promote the proximity of the AI warhead and Lys60, accounting for the largest percentage of the labeling reaction. Asp139 is involved in these immobilizations, and the LDAI reagent can also form intramolecular hydrogen bond interactions that help to constrain it for the reaction to take place.

Inspired by the interactions observed between the protein and the LDAI reagent in our MD simulations, Arg-, Asp-, and Glu-assisted mechanisms were incorporated into the QM reaction mechanisms, resulting in reduced energy barriers. The highest energy barriers for Argand Asp-assisted mechanisms of Lys labeling are 10.5 and 10.7 kcal/mol, respectively, and 11.4 and 11.9 kcal/mol for Glu- and Asp-assisted mechanisms of Ser labeling. As in aqueous solution, it is noted that there is a preference for Lys over Ser; however, consistent with the experimental reports, Glu and Asp assistance clearly distinguishes the Ser labeling of the reagent hydrolysis. Contrasting the energetics with the reported labeling percentages exposes an inconsistency regarding Ser labeling and the corresponding energy barriers, which we propose may stem from a lower likelihood of productive encounters between the warhead and Ser14 than between the warhead and Ser140.

The participation of Ser140 in LDAI reagent immobilization for Lys labeling supports our hypothesis that Ser140 contributes to Lys labeling, providing at least part of the explanation for the lack of labeling of GluA1, which carries the Ser140Ala substitution. The Ala residue is unable to form the double hydrogen bond that Ser140 forms with CAM2, directly implicating Ser140 in the labeling mechanism. However, this does not eliminate the possible role of Arg172 in Lys labeling and therefore does not fully explain the complete absence of labeling in GluA1. Based on our hydrogen bonding analysis, we speculate that in the absence of Ser140, Asp139 would form a more stable salt bridge with Arg172, competing with the roles of both Arg172 and Asp139 in Lys labeling. Interestingly, Arg, Asp, and Ser residues are also found in the vicinity of the labeled Lys in GABA_A_R,^57^ suggesting that the mechanistic motifs identified here may extend beyond AMPAR to other LDAI labeling targets. Depending on its chemical nature and linker length, the probe fragment may engage in interactions with the protein that affect the frequency of productive warhead–nucleophile encounters, providing a mechanistic explanation for the screening-like effects previously observed in labeling experiments.^2,58^

Based on our molecular modeling, we propose the following design principles to guide the rational development of future LDAI reagents, with potential generalizability to other warhead chemistries: (i) The spacing between the ligand fragment and the warhead must be sufficient for the warhead to effectively access a suitable nucleophile within the binding pocket. (ii) The target nucleophile should be flanked by neighboring residues, such as Arg, Asp, or Glu for LDAI chemistry, capable of facilitating the reaction through oxyanion stabilization or nucleophilic activation. (iii) The hydrogen-bond donors and acceptors on the LDAI reagent should be configured to interact cooperatively with complementary protein hydrogen-bond partners, such as Thr and Ser residues, maximizing reagent immobilization and warhead pre-organization. Together, these principles provide a structure-based framework for the rational optimization of LDAI reagents with improved selectivity, efficiency, and broader applicability across labeling targets.

## Methods

The following sections describe the computational protocols employed at the QM and MD levels, including the model systems, level of theory, simulation parameters, and analysis methods used to characterize the LDAI labeling mechanism and the GluA2–CAM2 interaction landscape.

### p***K*_a_** calculations

The protonation state of the acyl imidazole group in CAM2 directly affects the QM energy barriers of the labeling reaction, making knowledge of its p*K*_a_ essential for correctly weighting the Im^0^ and ImH^+^ contributions under experimental conditions. Since this value has not been reported, we estimated the p*K*_a_ of methyl 4-methyl-1H-imidazole-1-carboxylate, our cropped model of CAM2, with reference to a selection of similar compounds with reported p*K*_a_ values in the IUPAC Digitized p*K*_a_ Dataset^59^ (Table 4). We calculated p*K*_a_*^calc^* values for each compound following the deprotonation reaction

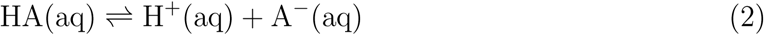

**Table 4:**
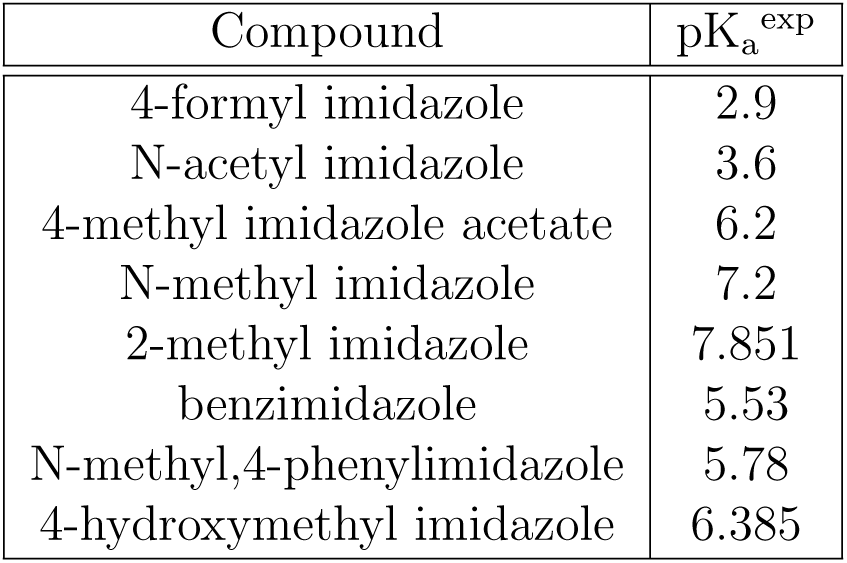
Imidazole derivates compound and their reported pK_a_ values selected to estimate the pK_a_ value of CAM2 by a linear regression.

and its associated ΔG^0^_deprot_ expression

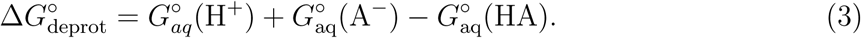

The standard aqueous Gibbs free energies of the conjugate base and acid, *G*_aq_*^◦^* (*A^−^*) and *G*_aq_*^◦^* (*HA*), were taken from the DFT calculations, while the standard aqueous Gibbs free energy of the proton, *G^◦^*_aq_ (*H*^+^), was determined using the following equation:

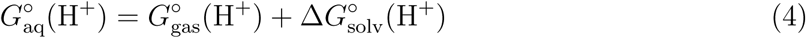

where *G^◦^*_gas_ is the Gibbs free energy of the proton in vacuum, and Δ*G^◦^*_solv_ is the solvation free energy, which are *−*6.28 kcal/mol and *−*265.63 kcal/mol, respectively.^60,61^ Using the following equation

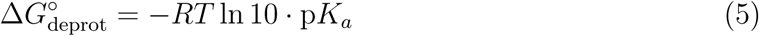

the p*K*^calc^*_a_* can be calculated as

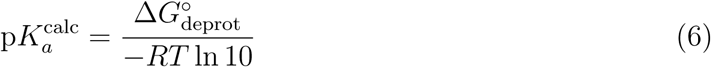

p*K*^calc^*_a_* values were obtained at *T* =298 K, and a linear regression of p*K*^exp^*_a_* versus p*K*^calc^*_a_* was performed, as depicted in Figure S2. The resulting y-intercept *C*_0_ and slope *C*_1_ were then used to estimate p*K*^fit^*_a_* for CAM2, following the expression

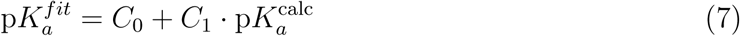

All geometry optimizations and vibrational frequency calculations were carried out using the Gaussian 16 program package,^62^ within the framework of density functional theory (DFT).^63,64^ Geometries and frequencies were obtained at the B3LYP/def2-SVPD level of theory,^35,37,65^ and the optimized structures were subsequently employed for single-point energy calculations at the def2-TZVPD level^37,66^ using the DSD-PBEP86 functional.^67^ All of the calculations include Grimme’s D3(BJ) empirical dispersion correction ^38,39^ and were performed with the Polarizable Continuum Model (PCM) to describe solvation in water.^40^

### Energy profile calculations

All geometry optimizations and vibrational frequency calculations were carried out using the CAM-B3LYP/def2-svp level of theory,^35–37^ including Grimme’s D3(BJ) empirical dispersion correction,^38,39^ and implicit water solvation with the Polarizable Continuum Model (PCM).^40^ For each proposed mechanism, the reference was its respective reactive complex. For the activation fees, CAM2 p*K*^fit^ of 4.33, with Lys p*K_a_* of 10.8 and a pH of 7 were employed.

### MD simulations

#### GluA2(S1S2J)-PFQX system

##### System preparation

The dimeric crystallizable construct of AMPAR GluA2, ^49^ GluA2(S1S2J) was generated with the Modeller software,^47,48^ using the X-ray structure of the GluA2 tetramer in complex with the ZK 200775 antagonist (PDB ID: 3KG2)^46^ as a template and the sequence reported in the experiments of Wakayama et al..^4^ Starting from the resulting GluA2(S1S2J)-ZK 200775 dimer, the quinoxaline scaffold of the PFQX antagonist was aligned to the ZK 200775 with Open Babel^68^ and replaced to create the GluA2(S1S2J)PFQX dimer. Topology files were prepared with the AmberTools23^69^ tleap module using the force field of proteins ff14SB ^70^ and the TIP3P^71^ water model. PFQX was parametrized using Antechamber and the GAFF2^72,73^ force field. Disulfide bonds Cys206-Cys261 were defined, and the GluA2(S1S2J)-PFQX dimer was solvated in 20 °A water molecules in all directions, forming a rectangular box. Na^+^ and Clions were added to obtain 0.15 M NaCl.

##### Energy minimization and MD simulations

The energy of the system was minimized using 1000 steepest descent steps followed by 1000 conjugate gradient steps, applying cartesian positional restraints to the protein main chain and PFQX quinoxaline scaffold with a force constant of 25 kcal/mol/°A^2^. Full electrostatic forces were treated using the PME method,^74^ and van der Waals interactions were computed using a 10 °A non-bonded cutoff radius. All molecular dynamic simulations were run with a 2 fs time step and constrained hydrogen-containing bonds using the SHAKE algorithm.^75,76^ Langevin dynamics^77^ was used for temperature control with a collision frequency of 2 ps^-1^. The systems were heated by linearly varying the target temperature from 0 to 310 K for 80 ps at constant volume, maintaining it at 310 K for an additional 20 ps, restraingin the protein main chain and PFQX quinoxaline scaffold with a force constant of 25 kcal/mol/°A^2^. Next, a restricted NPT equilibration was carried out at 310 K and 1 bar of pressure for 5 ns, applying cartesian positional restraints to the protein main chain and PFQX quinoxaline scaffold with a force constant of 25 kcal/mol/°A^2^. Then five restraintless independent NPT simulations of 2 µ*s* each were carried out.

##### Analysis

We clustered the simulation data using the DBSCAN algorithm of Ester et al. ^78^ by using the top and bottom distances, defined as the monomer-monomer distance between Ser229(Cα) and Lys185(Cα), respectively. One frame every 500 ps was selected from the five replicates, accumulating 20000 frames. An epsilon distance of 0.65 °A between points for forming a cluster and a minimum of 100 points (0.5% of analyzed frames) to form a cluster were set, leading to the definition of clusters as C1 and C2, along with some non-classified points, as shown in Figure 3B.

#### GluA2(S1S2J)-CAM2 system

##### Systems preparation

The centroids of clusters C1 and C2 were employed as the initial protein conformations for the C1 and C2 sets of GluA2(S1S2J)-CAM2 dimerOk simulations. One CAM2 molecule was parametrized using Antechamber and the GAFF2^72,73^ force field, and it was solvated in a 30 °A water box on each axis with 0.15 M NaCl and underwent a standard MD protocol of energy minimization, heating, and a 1 ns NPT production at 310 K and 1 bar. Then, a 100 ns 2D metadynamics^79^ simulation using PLUMED^80^ was carried out with end-to-end distance and radius of gyration as collective variables, with sigma values of 1 °Aand gaussian heights of 0.5 kJ/mol to obtain 10000 diverse conformations of CAM2. For the C1 and C2 seeds, each monomer was paired with one CAM2 conformation by aligning the CAM2 conformers and the PFQX quinoxaline scaffold using Open Babel, ^68^ resulting in 40000 GluA2(S1S2J)-CAM2 monomer complexes. Each of these monomer complexes was then subjected to a single-point energy calculation in sander, employing the GB-Neck2^81^ implicit solvent model. For every monomer complex, the 10 lowest-energy GluA2(S1S2J)-CAM2 conformers were retained. From these 10, the three most distinct complexes, as determined by their *RMSD* values, were selected and combined, resulting in a total of nine dimers in each set. Finally, each GluA2(S1S2J)-CAM2 dimer was embedded in a rectangular box of water extending 20 °A beyond the protein in all directions. Na^+^ and Clions were subsequently added to reach a final NaCl concentration of 0.15 M.

##### Energy minimization and MD simulations

Each resulting GluA2(S1S2J)-CAM2 dimer system was first subjected to energy minimization, consisting of 1000 steps of steepest descent followed by 1000 steps of conjugate gradient, while applying cartesian positional restraints to the protein backbone with a force constant of 25 kcal/mol/°A^2^. Electrostatic forces were treated with PME,^74^ and van der Waals interactions used a 10 °A non-bonded cutoff. All MD trajectories were computed with a 2 fs integration time step and SHAKE constraints on all bonds involving hydrogen atoms.^75,76^ Temperature was controlled by Langevin dynamics^77^ with a 2 ps*^−^*^1^ collision frequency. Systems were heated from 0 to 310 K over 80 ps at constant volume and held at 310 K for 20 ps, with the protein main chain restrained by a 25 kcal/mol/°A^2^ force constant. Next, three consecutive NPT equilibration steps were carried out at 310 K and 1 bar of pressure for 10 ns each, applying cartesian positional restraints to the protein main chain with force constants of 25, 10, and 5 kcal/mol/°A^2^, respectively. Finally, all restrictions were removed, and an NPT simulation of 200 ns at 310 K and 1 bar was performed.

##### Analysis

*RMSD*, distance, and hydrogen bond calculations were performed using cpptraj.^82^ Detailed analysis and graphs were produced using in-house Python ^83^ scripting. Structure images were generated using PyMOL.^84^ The standard criteria of cpptraj for hydrogen bond definition were employed, i.e., a cutoff distance of 3 °A between the donor (D) and acceptor (A) atoms, and an angle of at least 135° ^(^*^̸^* DHA).

## Acknowledgement

The authors acknowledge support from the IKUR Strategy (Nanoneuro) under the collaboration agreement between Ikerbasque Foundation and DIPC and the “Programa de Ayuda de Apoyo a los agentes de la Red Vasca de Ciencia, Tecnoloǵıa e Innovación acreditados en la categoŕıa de Centros de Investigacíon Básica y de Excelencia (Programa BERC)” from the Departamento de Ciencia, Universidades e Innovación del Gobierno Vasco and Centros Severo Ochoa AEI/CEX2024-0001491-S from the Spanish Ministerio de Ciencia e Innovación. DDS and XL receive support from the Basque Government through Project IT2067-26 and grant PID2024-158678NB-I00 funded by MICIU/AEI/10.13039/501100011033 and “ERDF A way of making Europe”.

## Supporting Information Available

The Supporting Information includes additional energy profiles and transition-state energy barriers, the linear regression used to determine the p*K_a_* of the imidazole ring, and complementary analyses of distances, RMSD, and hydrogen bonding in the MD trajectories. Initial coordinates as well as input/output files are available at https://doi.org/10.5281/zenodo.22140043.

## TOC Graphic

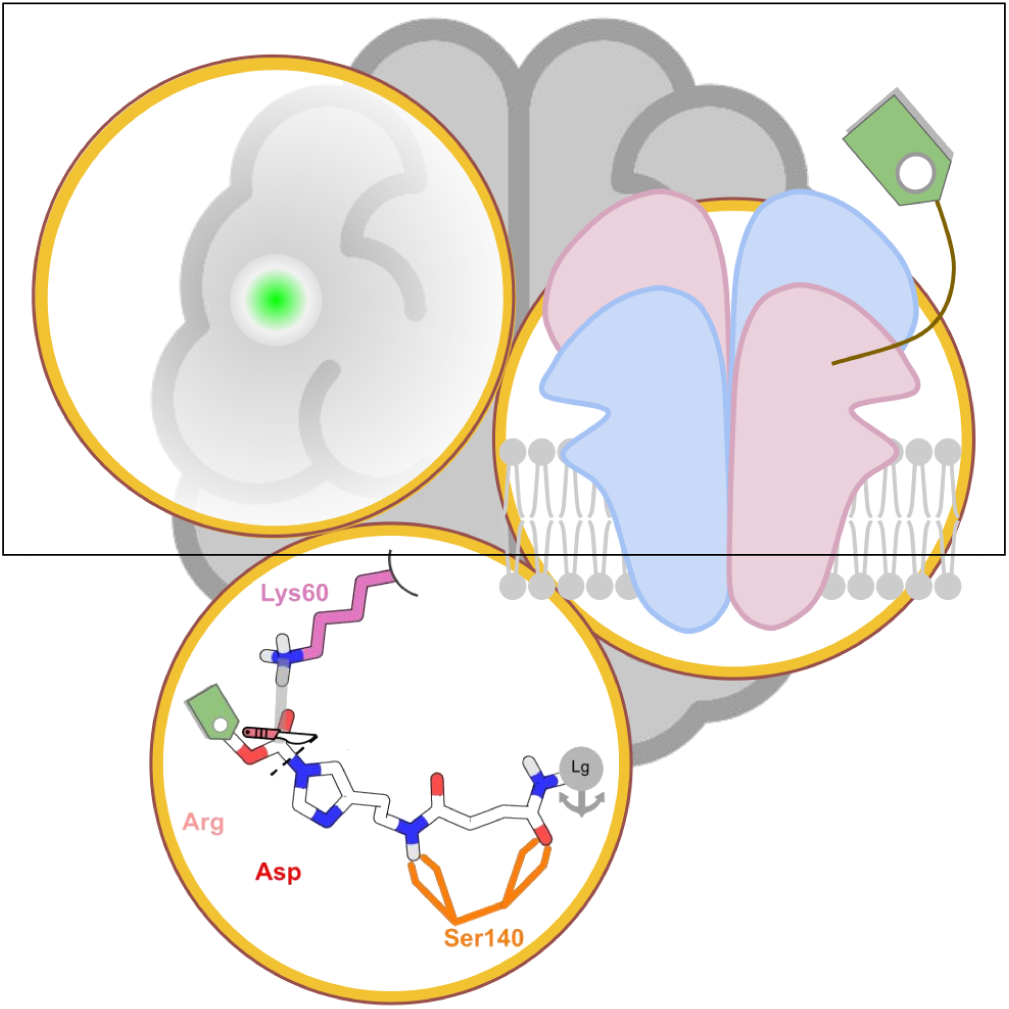

