## Supporting Information for "Molecular basis of AMPA receptor labeling by ligand-directed acyl imidazole chemistry in living neurons"

### Supporting Information: Mechanism of AMPA receptor labelling by ligand-directed acyl imidazole chemistry

Table S1: Sequence of the GluA2(S1S2J), the LBD crystallizable construct used in our simulations, identical to that employed to identify the labeled residues. Ser14, Ser140, and Lys60 are highlighted in green and blue, respectively. Residues belonging to the junction sequence are shown in gray. Numbering in parentheses refers to positions in the full-length protein.

```
   10(420)         20         30         40         50    60(470)
GANKTVVVTT ILESPYVMMK KNHEMLEGNE RYEGYCVDLA AEIAKHCGFK YKLTIVGDGK
   70(480)         80         90        100        110    120(653)
YGARDADTKI WNGMVGELVY GKADIAIAPL TITLVREEVI DFSKPFMSLG ISIMIKKGTP
  130(663)        140        150        160        170    180(713)
IESAEDLSKQ TEIAYGTLDS GSTKEFFRRS KIAVFDKMWT YMRSAEPSVF VRTTAEGVAR
  190(723)        200        210        220        230    240(773)
VRKSKGKYAY LLESTMNEYI EQRKPCDTMK VGGNLDSKGY GIATPKGSSL GNAVNLAVLK
  250(793)        260
LNEQGLLDKL KNKWWYDKGE CGS
```

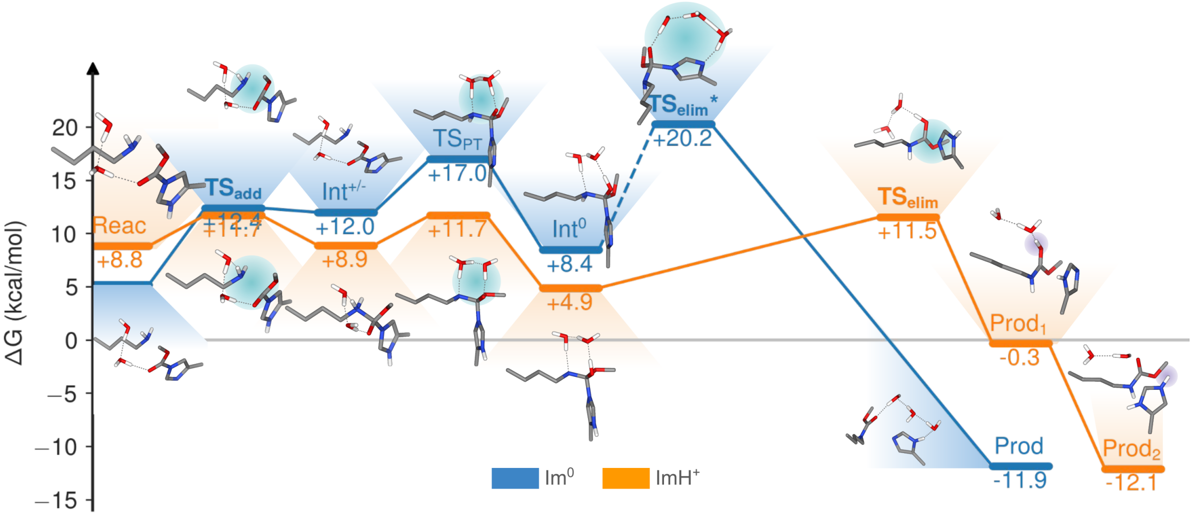


Figure S1: Energy profile of Lys labeling with both Im ring protonation states. $TS_{elim}$* indicates the energy barrier of the precursor step that directly triggers the subsequent elimination.

Table S2: Energy barriers of transition states for Lys labeling modifying the Im protonation state and number of explicit water molecules. With bold style is highlighted the higher energy barrier with $\Delta G_{prot}$ not added yet and gray fill indicate the lowest energy path between both elimination pathways. The numbers (1), (2), and (3) indicate the sequential order of the mechanistic steps.

<table>
<tr><th rowspan="2">System</th><th rowspan="2">$\Delta G_{prot}$</th><th rowspan="2">$TS_{add}$ (1)</th><th colspan="2">$TS_{elim}$ (2)</th></tr>
<tr><th>$TS_{PT}$(2)</th><th>$TS_{elim}$(3)</th></tr>
<tr><td>Lys·CAM2($Im^0$)</td><td>+5.2</td><td>-</td><td>42.3</td><td>35.9</td></tr>
<tr><td rowspan="2">Lys·CAM2($ImH^+$)</td><td rowspan="2">+8.8</td><td rowspan="2">17.4</td><td colspan="2">20.3</td></tr>
<tr><td>41.7</td><td>19.5</td></tr>
<tr><td rowspan="2">Lys·CAM2($ImH^+$)·$H_2O$</td><td rowspan="2">+8.8</td><td rowspan="2">13.1</td><td colspan="2">14.2</td></tr>
<tr><td>16.0</td><td>10.9</td></tr>
<tr><td rowspan="2">Lys·CAM2($Im^0$)·$2H_2O$</td><td rowspan="2">+5.2</td><td rowspan="2">12.4</td><td colspan="2">-</td></tr>
<tr><td>17.0</td><td>-</td></tr>
<tr><td rowspan="2">Lys·CAM2($ImH^+$)·$2H_2O$</td><td rowspan="2">+8.8</td><td rowspan="2">11.7</td><td colspan="2">12.8</td></tr>
<tr><td>11.7</td><td>11.5</td></tr>
<tr><td rowspan="2">Lys·CAM2($ImH^+$)·$3H_2O$</td><td rowspan="2">+8.8</td><td rowspan="2">11.7</td><td colspan="2">10.3</td></tr>
<tr><td>12.6</td><td>9.9</td></tr>
</table>

Table S3: Comparison of energy barriers of transition states (TS) for Lys and Ser labeling and CAM2 hydrolysis. With bold style is highlighted the higher energy barrier and gray fill indicate the lowest energy path between both elimination pathways. The numbers (1), (2), and (3) indicate the sequential order of the mechanistic steps. $\Delta G_{prot}$ values are added to the TS barriers. Geometry and frequency calculations were done with the CAM-B3LYP/def2-SVP, employing the optimized geometries to perform single-point energy evaluations at CAM-B3LYP/def2-TZVPD.

| System | $\Delta G_{prot}$ | $TS_{add}$ (1) / $TS_{add—PT}$ (1) | $TS_{elim}$ (2) | |
|---|---|---|---|---|
| | | | $TS_{PT}$(2) | $TS_{elim}$(3) |
| Lys·CAM2($ImH^+$)·2$H_2O$ | +8.8 | **13.5** | **13.5** | |
| | | | 17.1 | 15.1 |
| Ser·CAM2($ImH^+$)·2$H_2O$ | +3.6 | **23.7** | 16.0 | |
| CAM2($ImH^+$)·3$H_2O$ | +3.6 | **24.6** | 19.7 | |

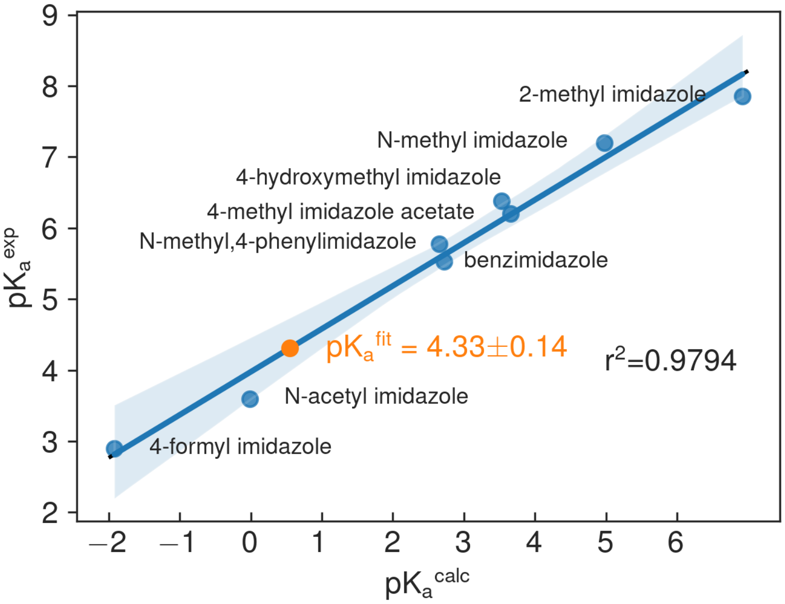


Figure S2: Linear regression for the determination of the $pK_a$ of the cropped-CAM2 Im ring. Geometry and frequency calculations were done with the B3LYP/def2-SVPD, employing the optimized geometries to perform single-point energy evaluations at DSD-PBEP86/def2-TZVPD.

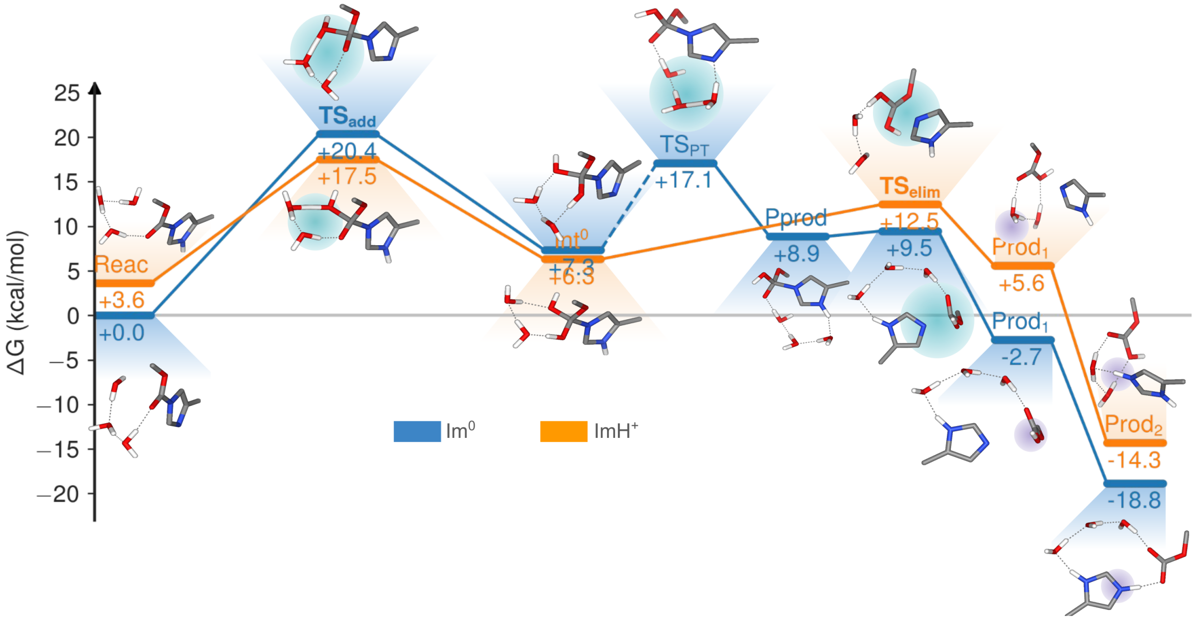


Figure S3: Energy profile of CAM2 hydrolysis with both Im ring protonation states.

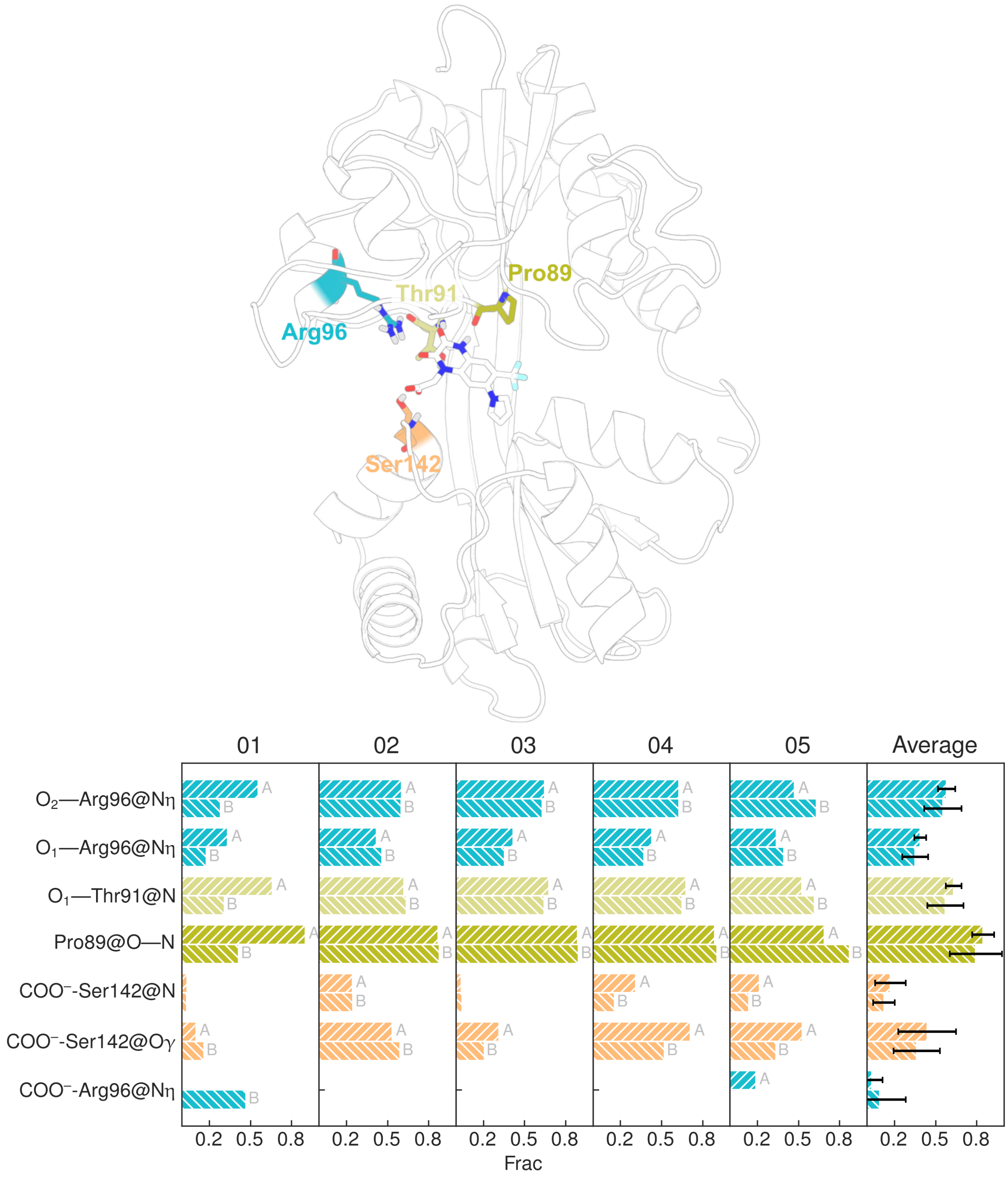


Figure S4: Fractions of hydrogen bond between PFQX antagonist and GluA2(S1S2J) protein for each monomer and replicate. Diagonal hatch (// and \\) distinguishes the monomers.

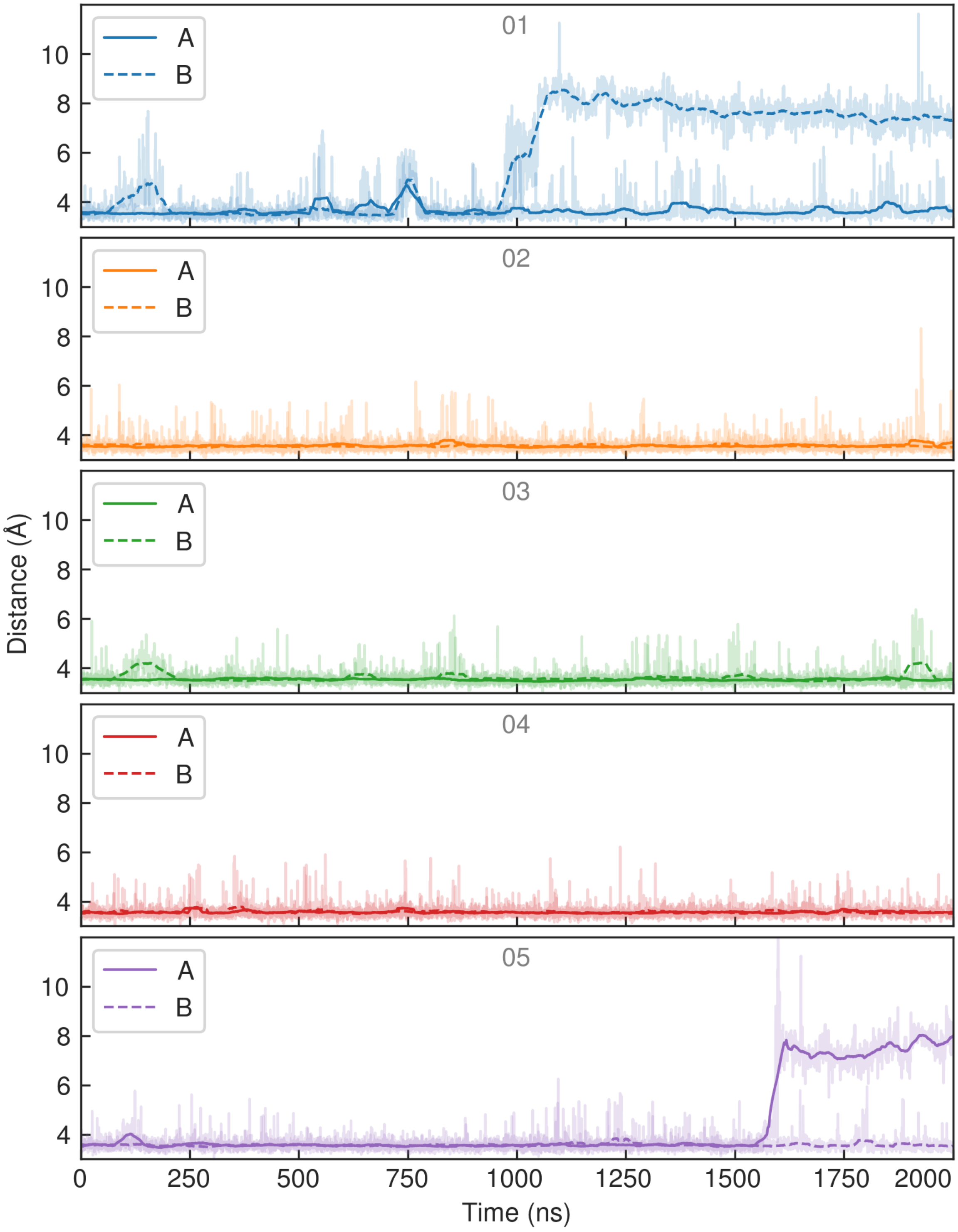


Figure S5: Time series of PFQX antagonist binding measured as the distance between GluA2(S1S2J) Arg96(C$\zeta$) and the center of mass of the quinoxalinedione oxygens for all five replicates and each monomer.

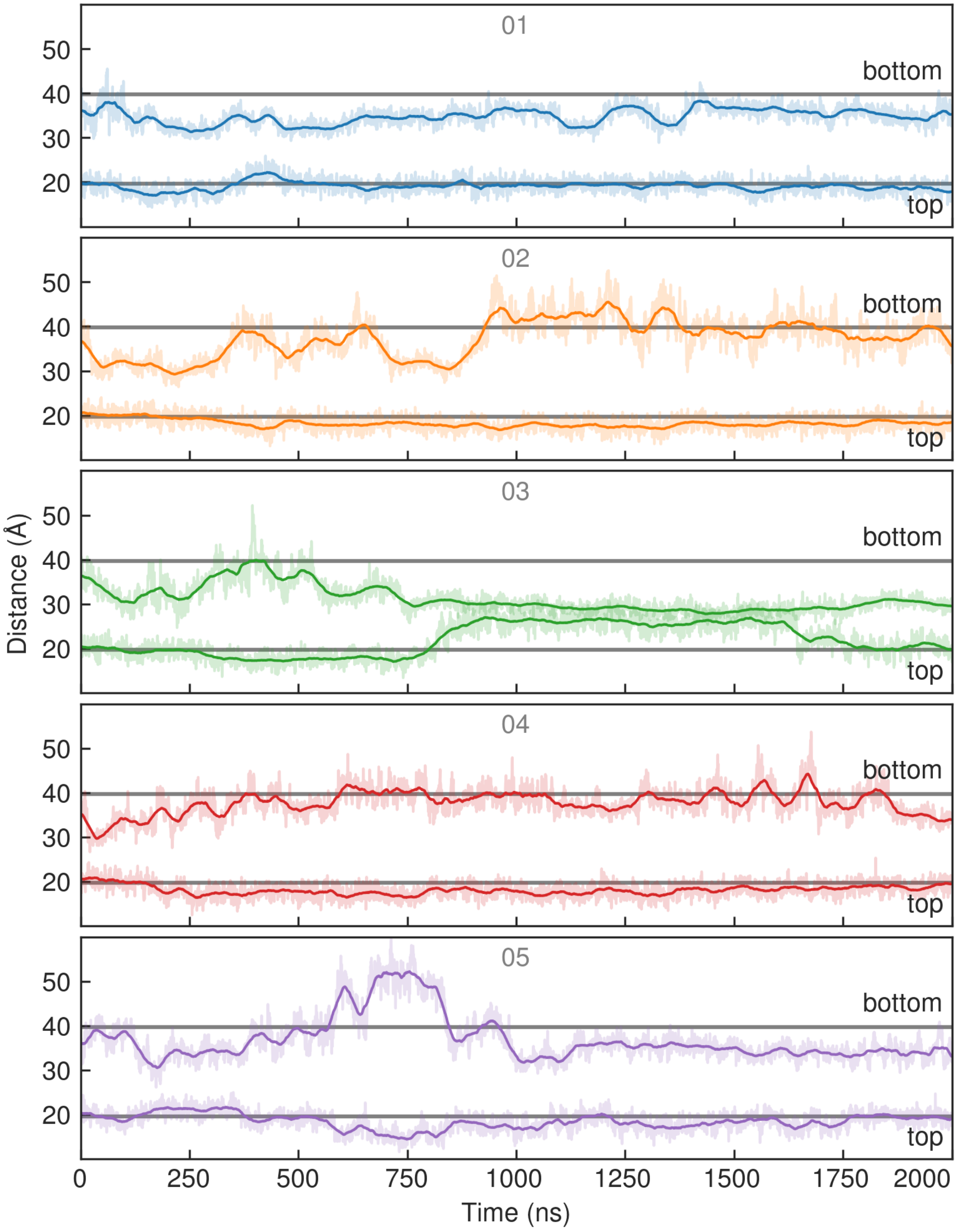


Figure S6: Time series of the top (upstream) and bottom (downstream) distances with respect the membrane in the whole protein. Top distance as the distances between monomer-monomer Ser229(C$\alpha$) and bottom distance as the distances between Lys185(C$\alpha$). Gray horizontal lines indicate the initial values of both distances.

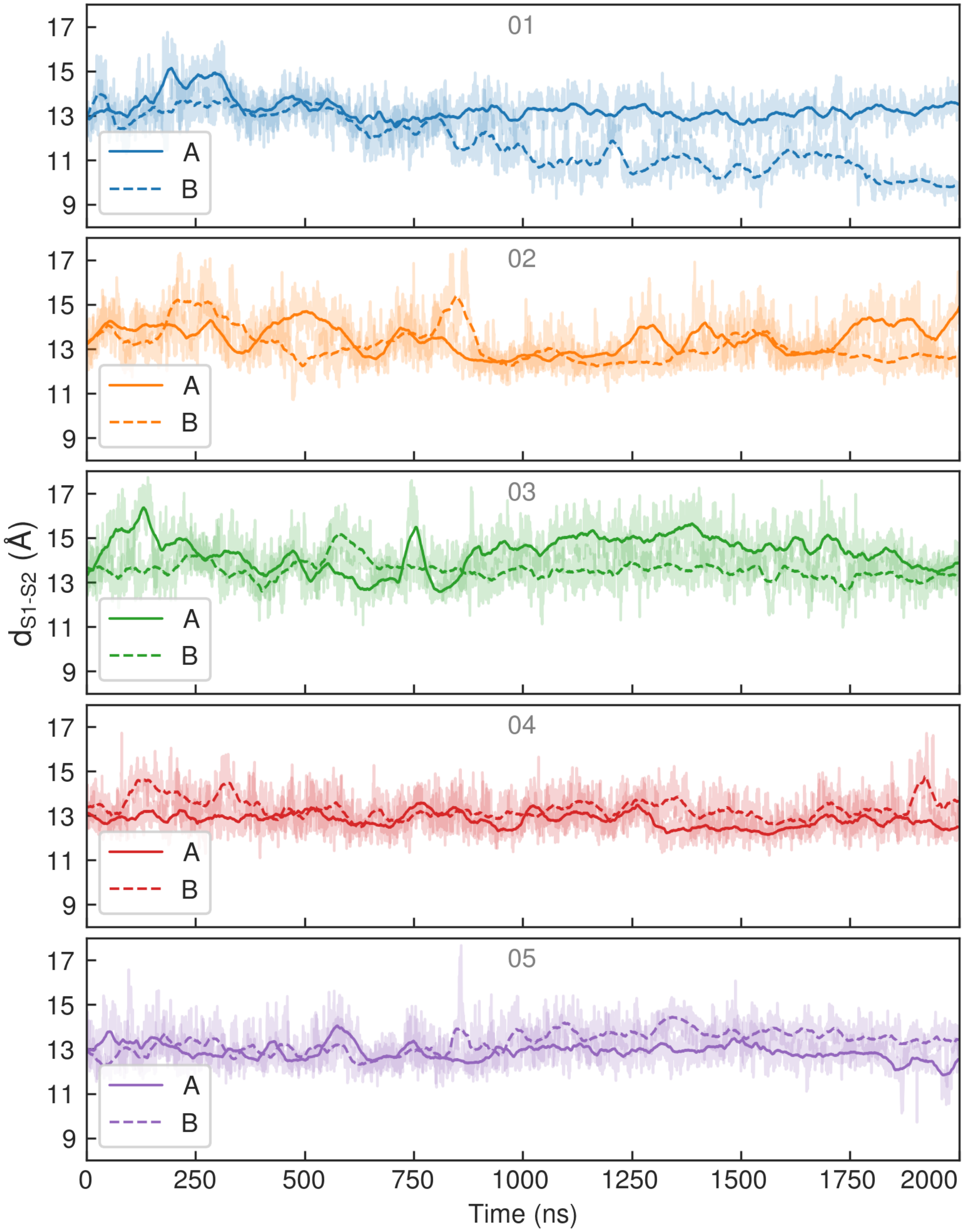


Figure S7: Time series of clamshell state measured as the distance between the center of mass of Leu90, Thr91 and Ile92 (S1 domain) and Ser142 and Thr143 (S2 domain).

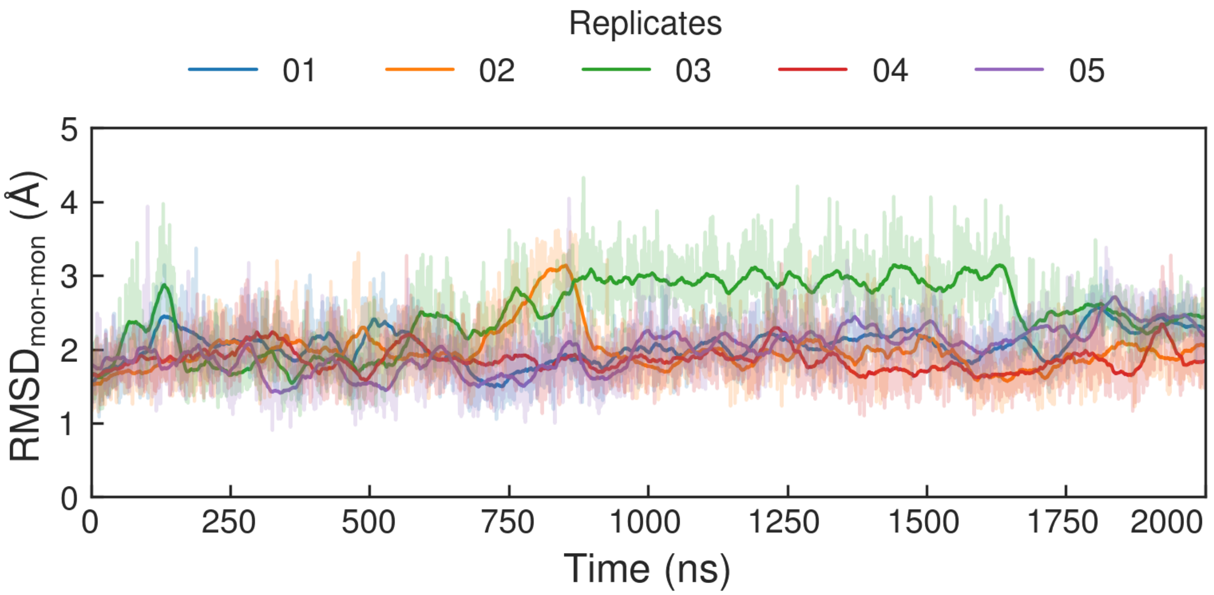


Figure S8: Root mean square deviation between monomer-monomer main chains for all five replicates.

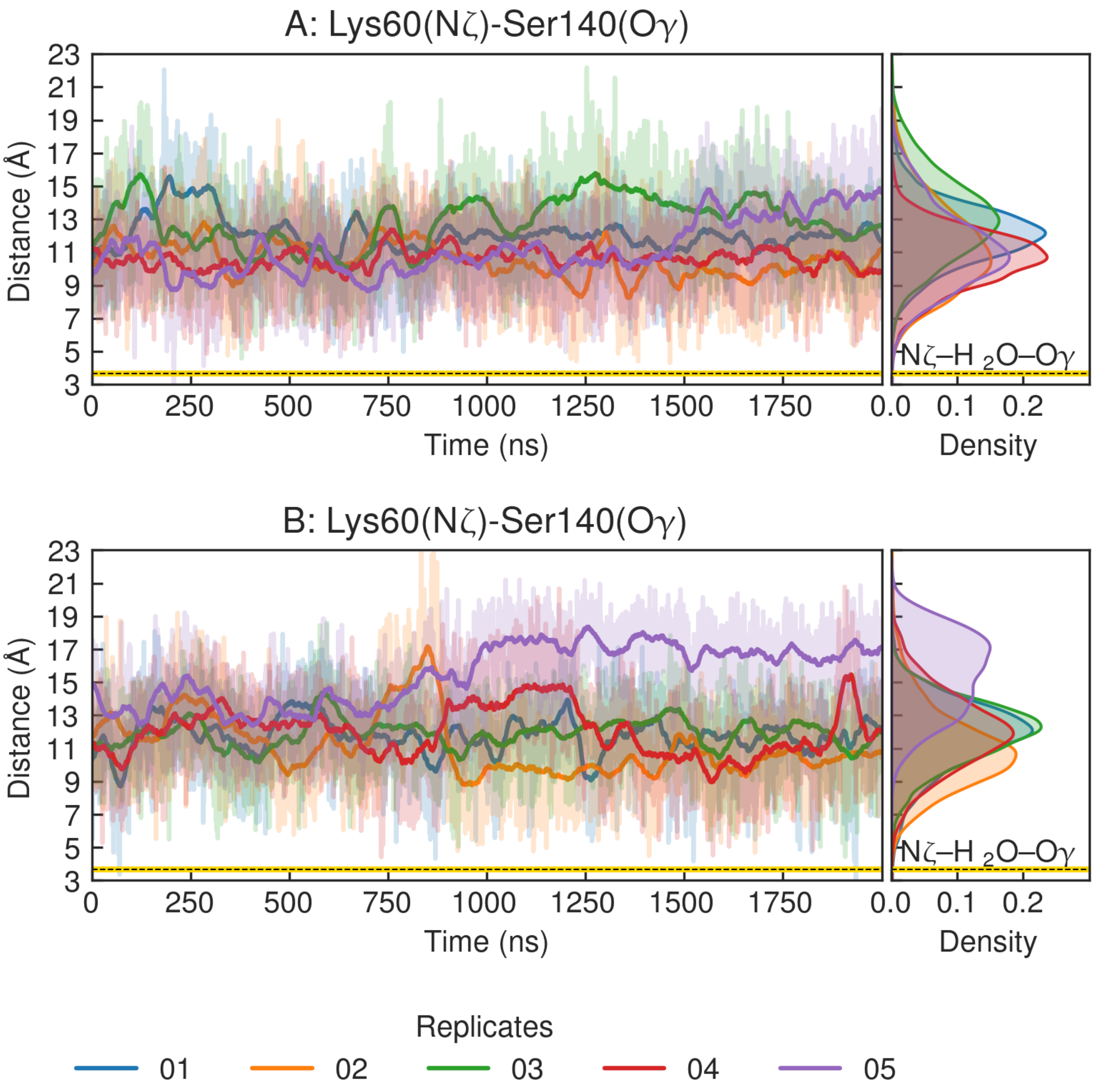


Figure S9: Time series and density distributions of the distance between the Lys60 and Ser140 side chains for each monomer in the five replicates. The black and yellow dashed line indicates the distance in the case of a one-water bridge.

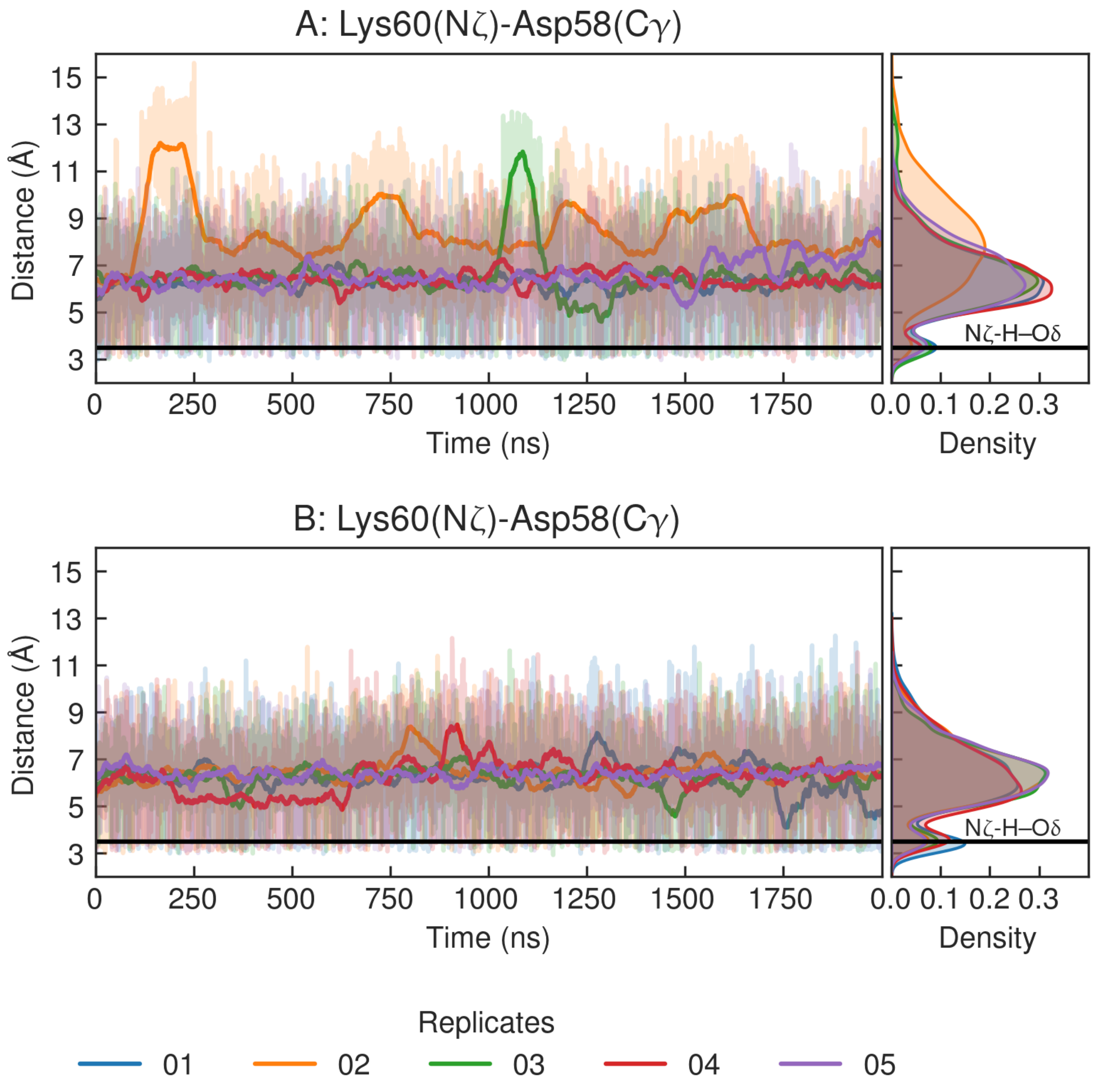


Figure S10: Time series and density distributions of the distance between the Lys60 and Asp58 side chains for each monomer in the five replicates. The black horizontal line indicates the distance in the case of a hydrogen bond.

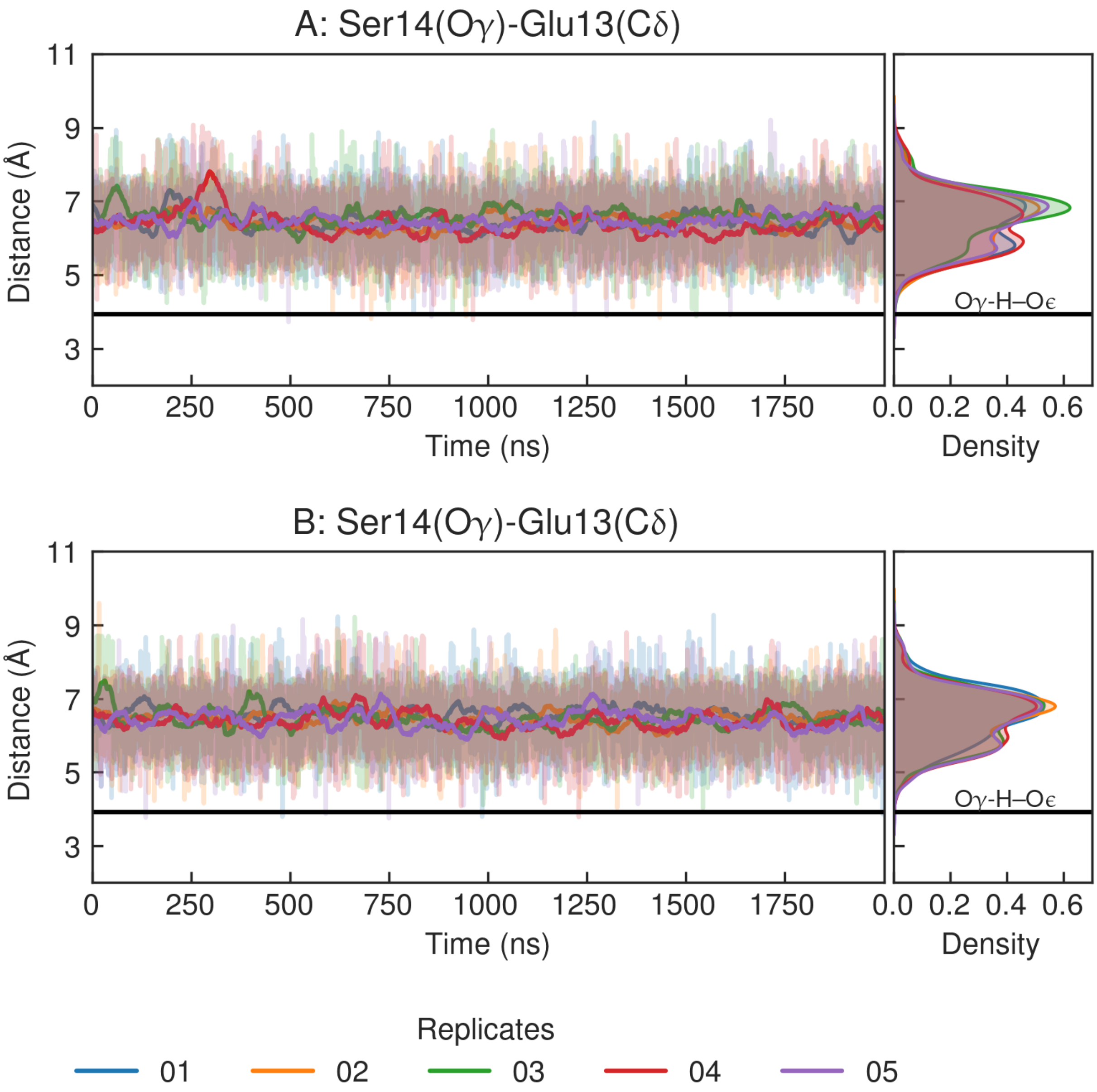


Figure S11: Time series and density distributions of the distance between the Ser14 and Glu13 side chains for each monomer in the five replicates. The black horizontal line indicates the distance in the case of a hydrogen bond.

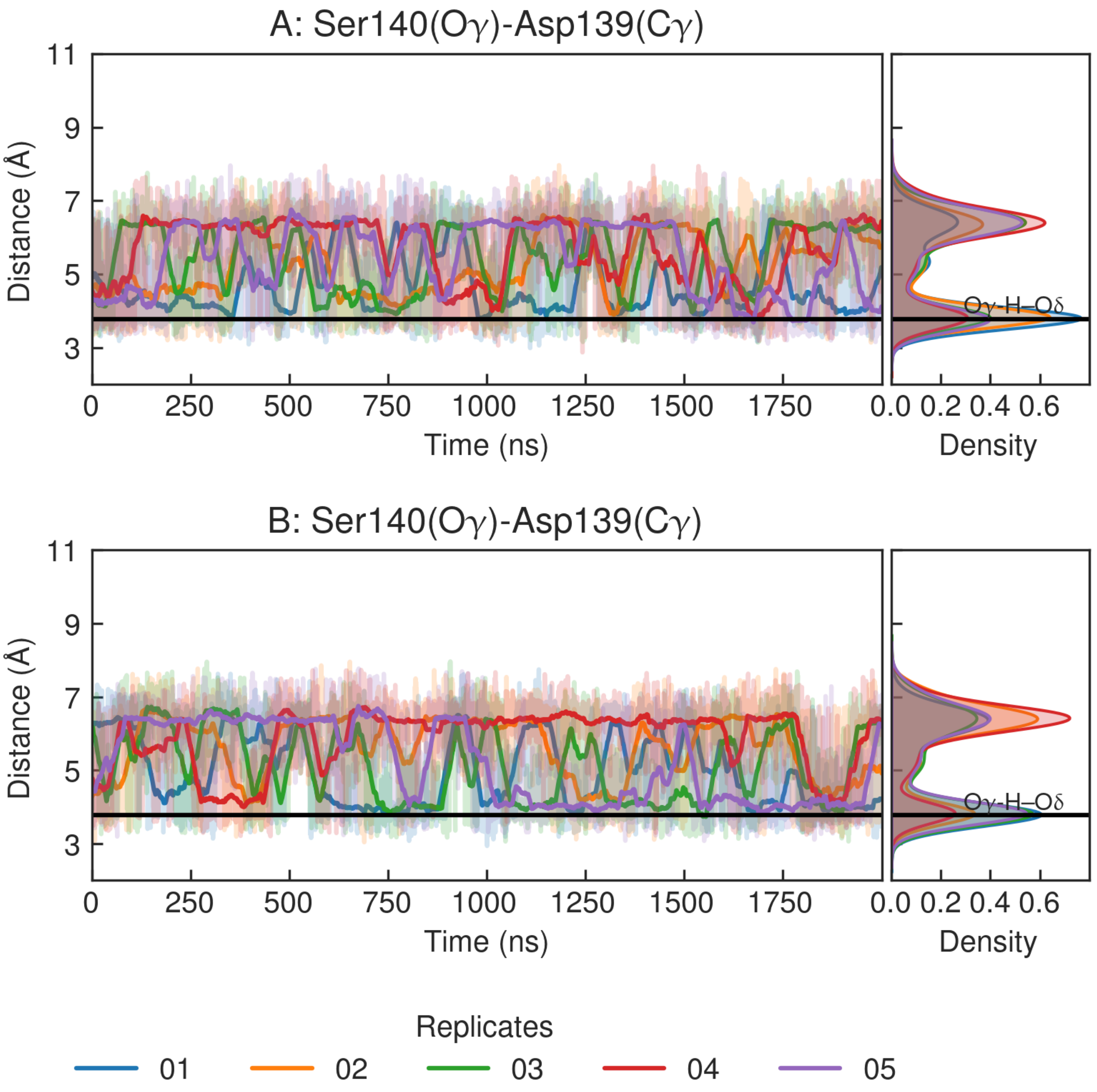


Figure S12: Time series and density distributions of the distance between the Ser140 and Asp139 side chains for each monomer in the five replicates. The black and yellow dashed line indicates the distance in the case of a hydrogen bond.

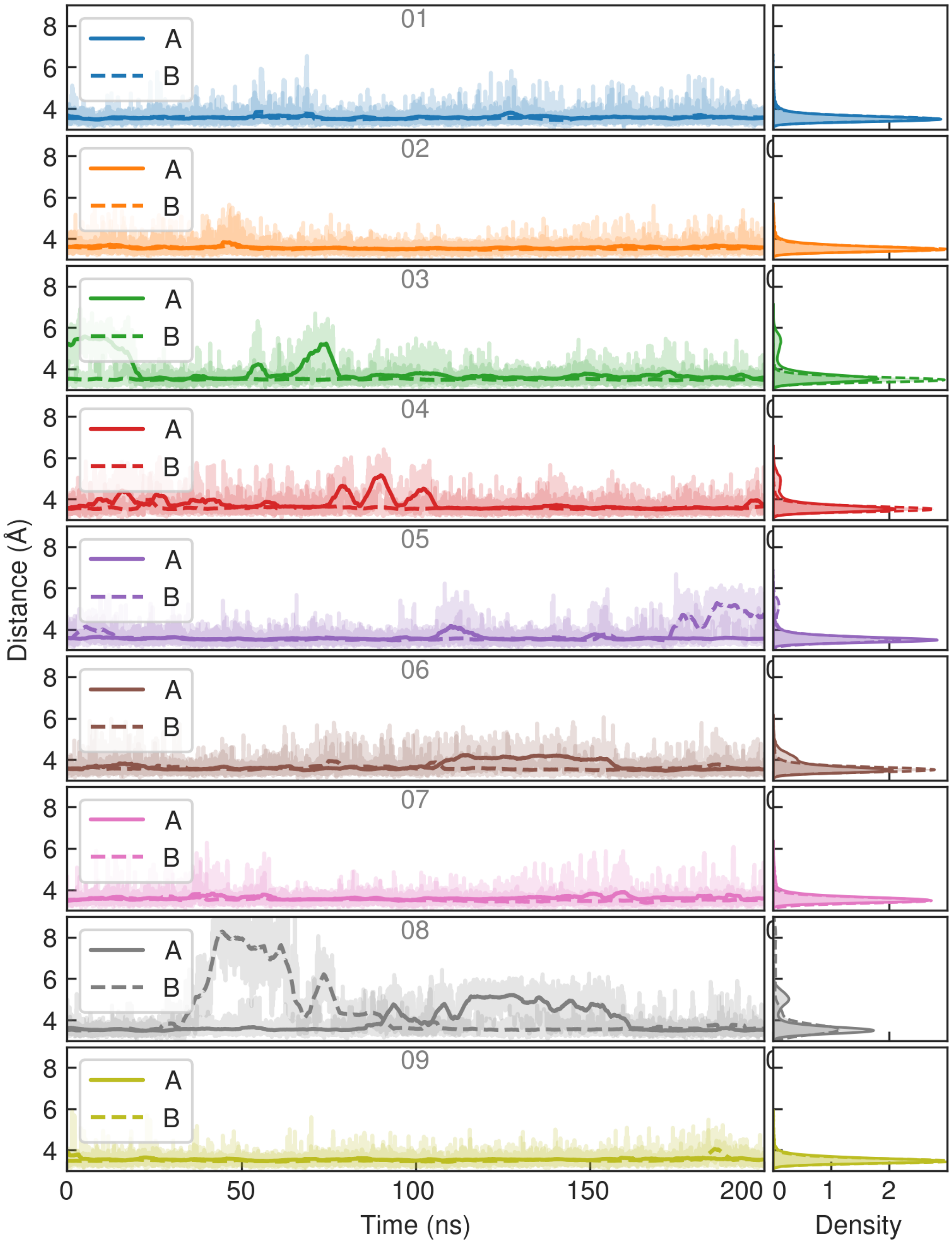


Figure S13: Time series of antagonist-moiety binding measured as the distance between Arg96(C$\zeta$) and the center of mass of the CAM2 quinoxalinedione oxygens for all nine replicates and each monomer of C1 set.

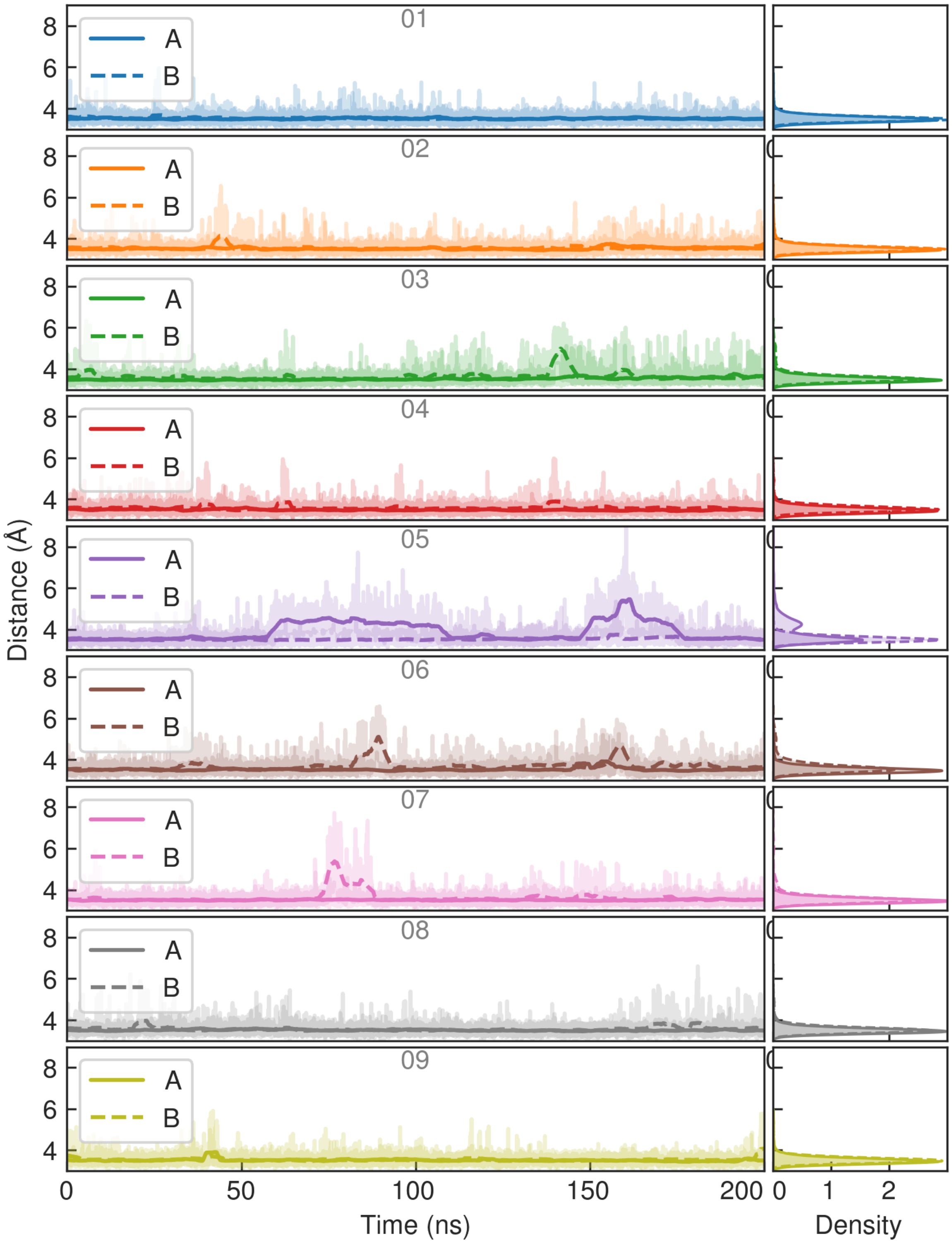


Figure S14: Time series of antagonist-moiety binding measured as the distance between GluA2(S1S2J) Arg96(Cζ) and the center of mass of the CAM2 quinoxalinedione oxygens for all nine replicates and each monomer of C2 set.

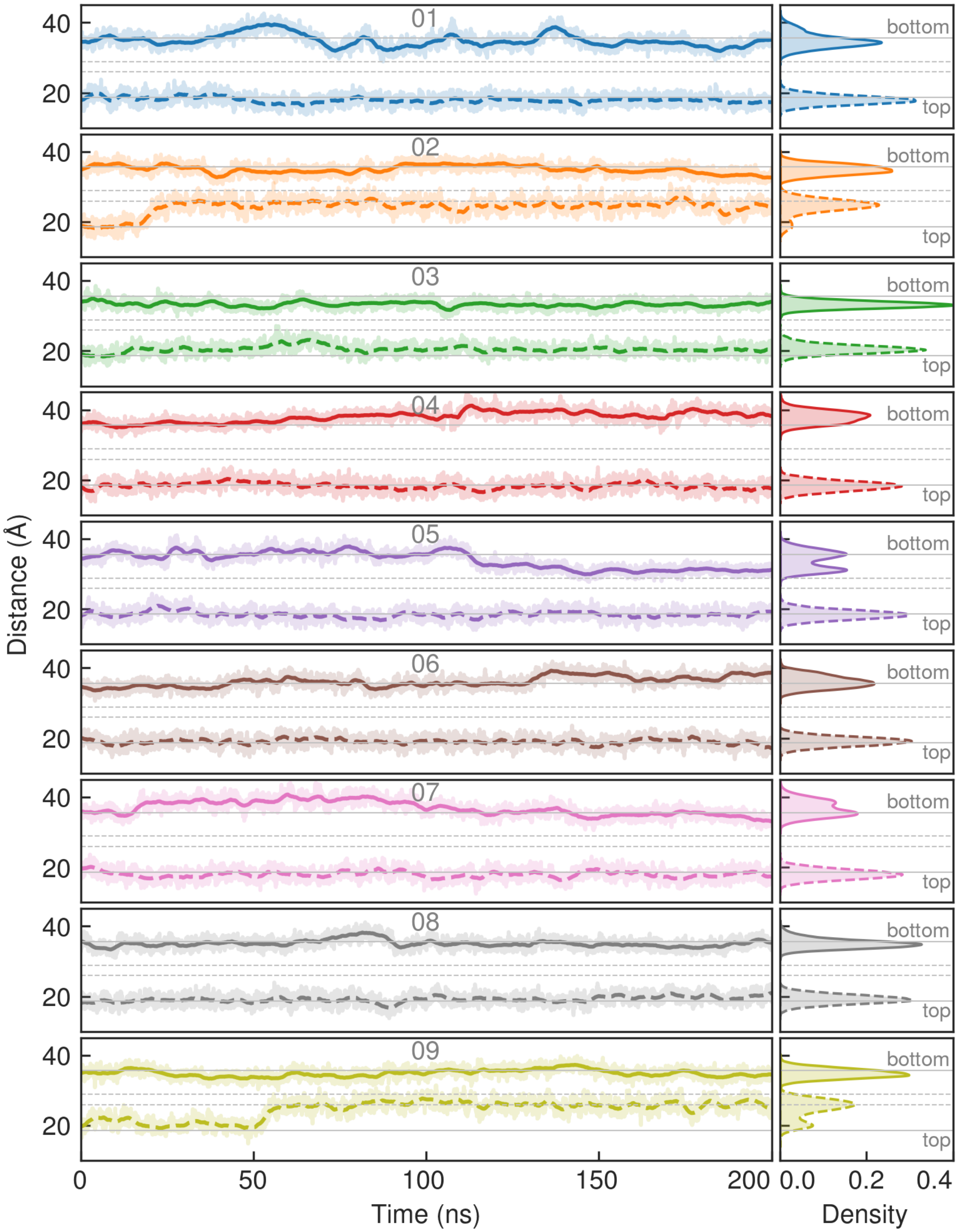


Figure S15: Time series of the top (upstream) and bottom (downstream) distances with respect the membrane in the whole protein. Top distance as the distances between monomer-monomer Ser229(C$\alpha$) and bottom distance as the distances between Lys185(C$\alpha$) of C1 set. Black horizontal lines show the initial distances for both sets; the current set is represented by the solid line.

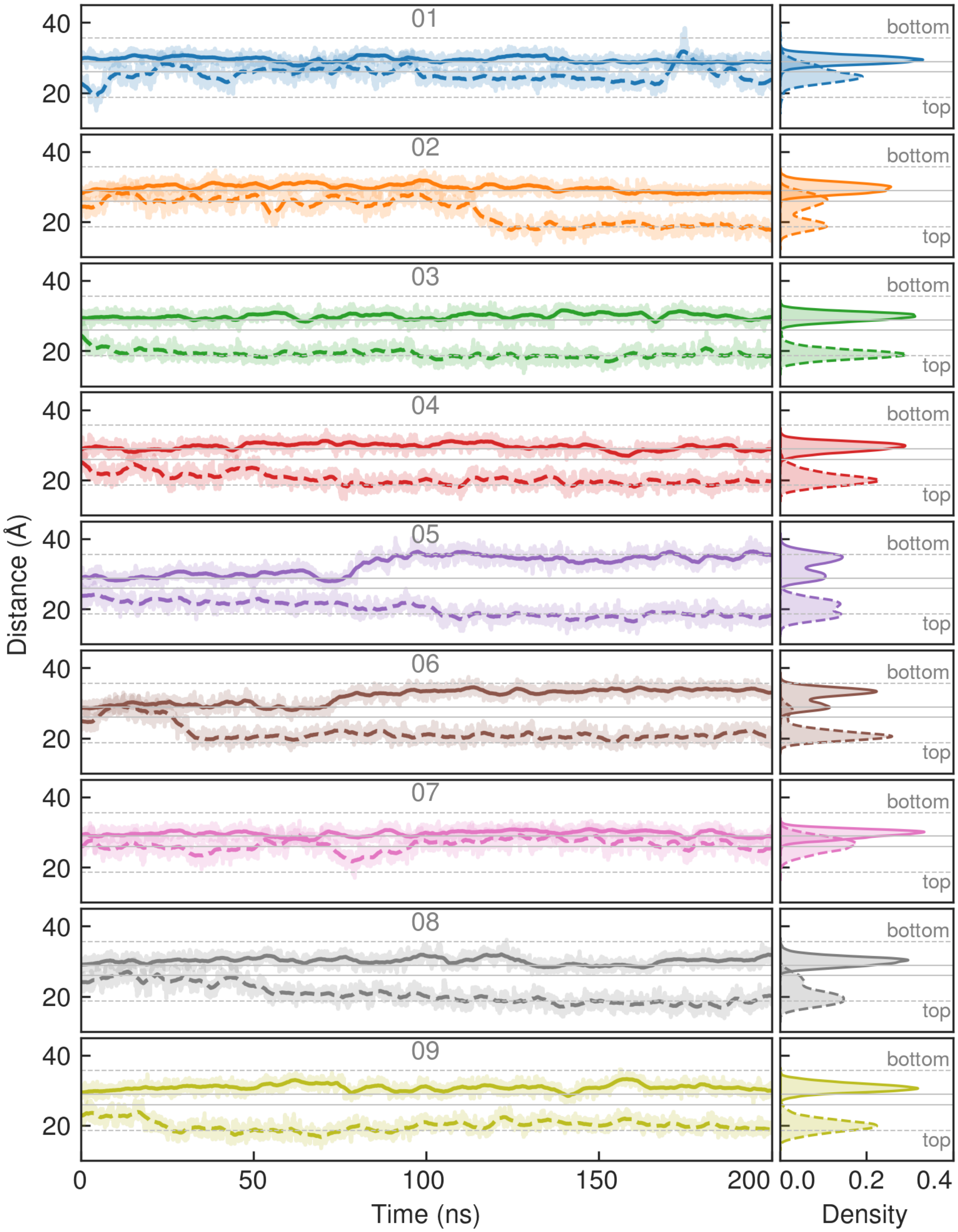


Figure S16: Time series of the top (upstream) and bottom (downstream) distances with respect the membrane in the whole protein. Top distance as the distances between monomer-monomer Ser229(C$\alpha$) and bottom distance as the distances between Lys185(C$\alpha$) of C2 set. Black horizontal lines show the initial distances for both sets; the current set is represented by the solid line.

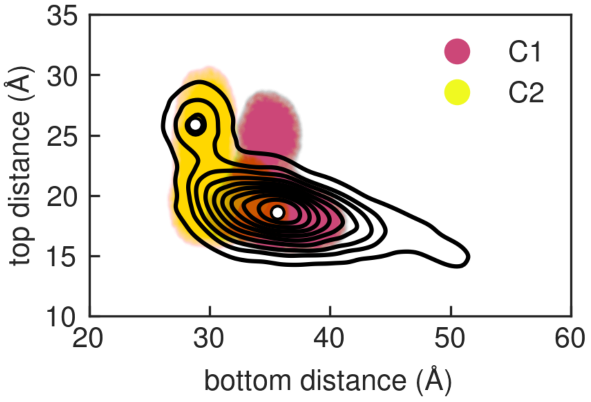


Figure S17: Top and bottom distances of C1 and C2 GluA2(S1S2J)-CAM2 sets onto the 2D clustering projection from GluA2(S1S2J)-PFQX simulations.

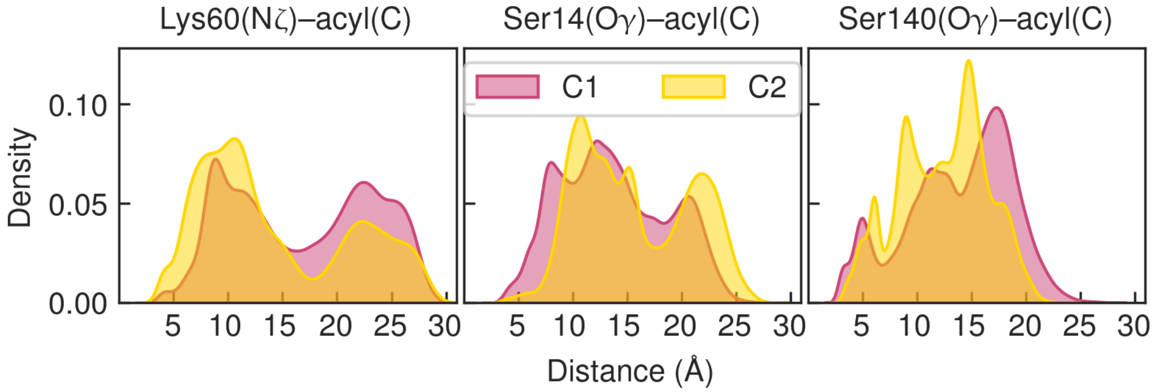


Figure S18: 1D distributions of the distances between each labeled nucleophile and the acyl carbon electrophile for both C1 and C2 sets.

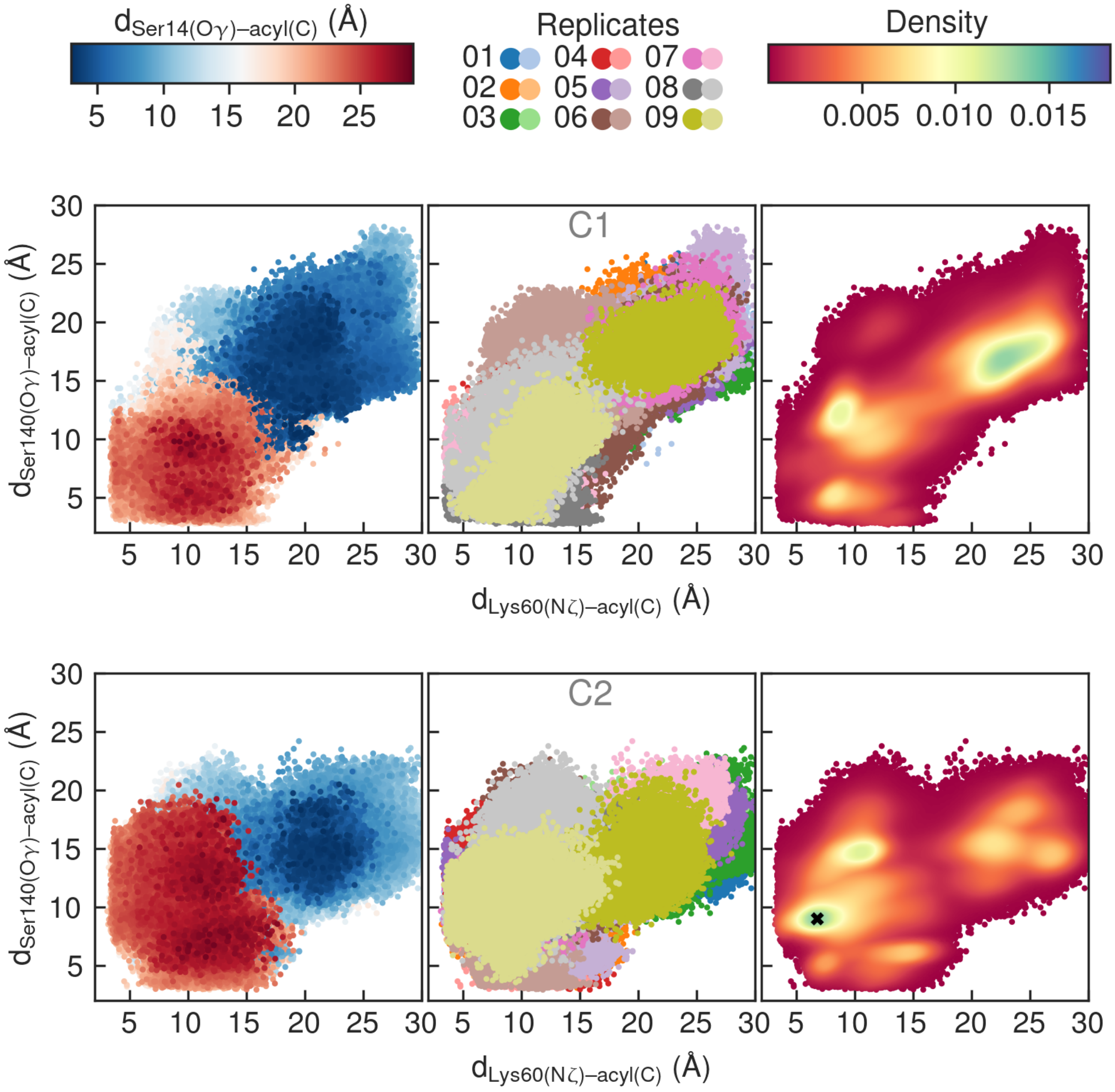


Figure S19: 2D distances distributions between CAM2 acyl cabron and Lys60(N$\zeta$) and Ser140(O$\gamma$) for all nine replicates of C1(above) and C2(below) sets. Same data is plotted from left to right but employing the color to indicate the respective distance with Ser14(O$\gamma$), the replicates and monomers populations and, the data density. × represents a high-density population present in set C2 but not in C1, identified as a geometry corresponding to a possible critical point where Ser140 is positioned to facilitate the labeling of Lys60.

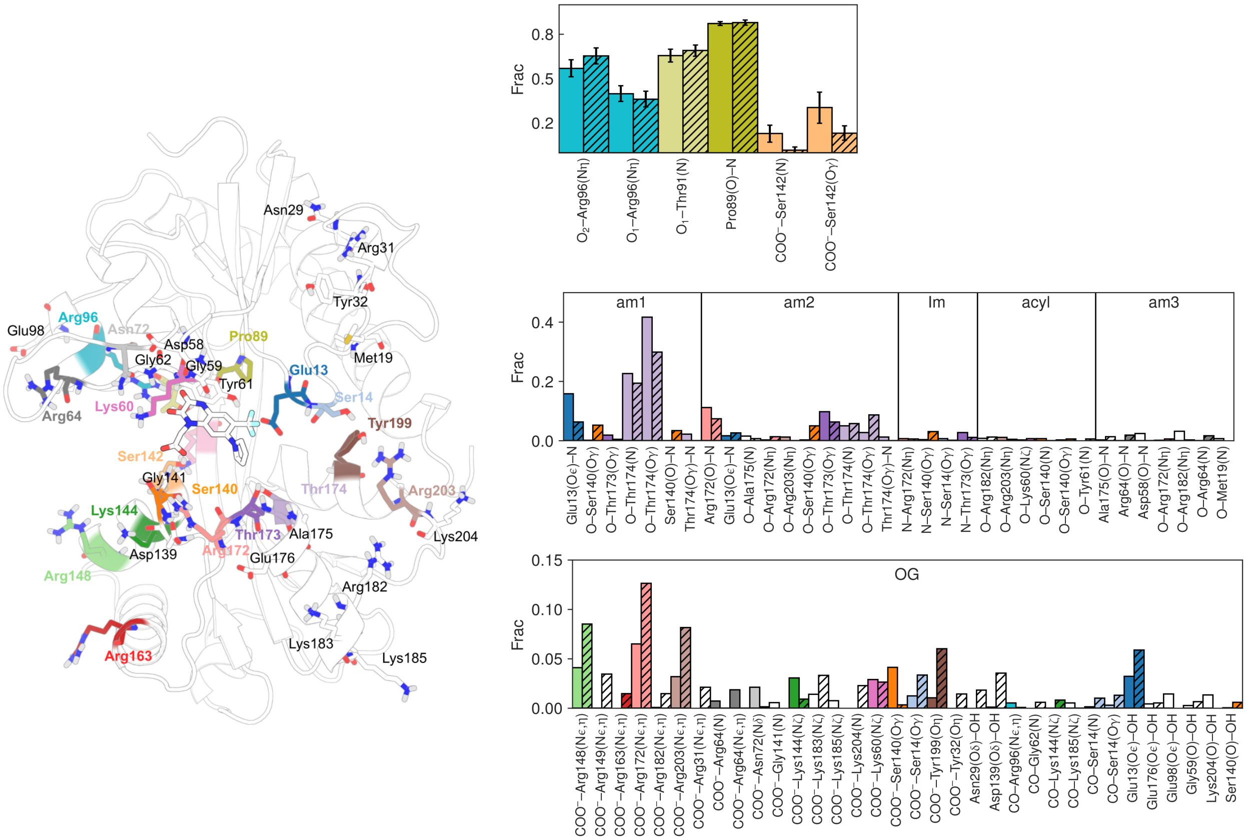

Figure S20: Barplots of fractions of hydrogen bond interactions of CAM2 with GluA2(S1S2J) protein. Graph is separated in three parts of CAM2, from above to below: the antagonist fragment (QX), the middle that include amide groups (am1-3) and reactive acyl and, Im groups and the oregon green (OG) fragment. C2 is distinguished from C1 by diagonal hatching (//).

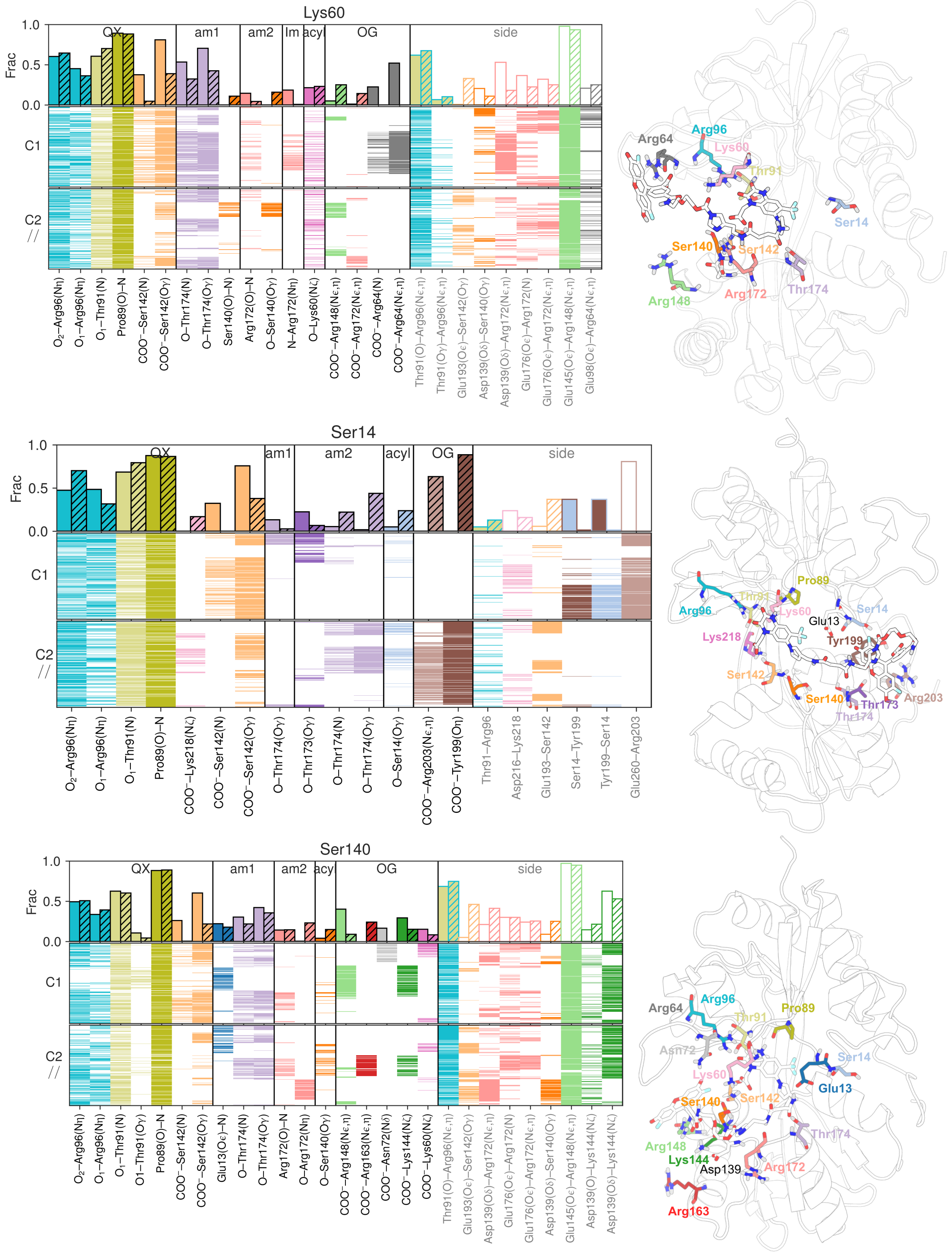


Figure S21: Barplots of fractions and presences of hydrogen bonds between GluA2(S1S2J) and CAM2 when each nucleophile Lys(Nζ), Ser14(Oγ) and Ser140(Oγ) are at least 5 Å from the CAM2 acyl carbon. Hydrogen bond interactions are ordered according CAM2 atoms from antagonist fragment (QX) to Oregon Green (OG). CAM2 amide groups are indicate as am1-3. Side hydrogen bond interactions involving residues that interact with CAM2 are also shown.

Table S4: Energy barriers of Lys labeling assisted by protein backbone and Ser residue. The numbers (1) and (2) indicate the sequential order of the mechanistic steps.

| System | $\Delta G_{prot}$ | $TS_{add}$ (1) | $TS_{elim}$ (2) | $TS_{max}$ | $Int^0$ |
|---|---|---|---|---|---|
| Lys·CAM2($ImH^+$)·$H_2O$· –NH–acyl(O)–HN– | +8.83 | 13.0 | **14.8** | 14.8 | 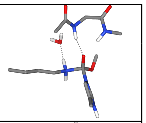 |
| Lys·CAM2($ImH^+$)· –CO–Lys($NH_2^+$), acyl(O)–HN– | +8.83 | **15.0** | 14.5 | 14.97 | 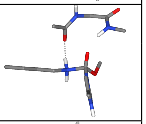 |
| Lys·CAM2($ImH^+$)·Ser | +8.83 | **14.2** | 5.26 | 14.1 | 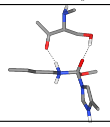 |

Table S5: Energy barriers of transition states for Lys labeling assisted by Arg and Asp. References are each respective reactive complex. Units are in kcal/mol. The gray fill highlights the lowest energy path, if applicable, and the bold formatting indicates the highest energy barrier before the activation fee. The numbers (1), (2), and (3) indicate the sequential order of the mechanistic steps. * denotes the energy barrier associated with transfer of the Lys proton to the Asp that leads to the elimination. Geometry and frequency calculations were done with the CAM-B3LYP/def2-SVP, employing the optimized geometries to perform single-point energy evaluations at CAM-B3LYP/def2-TZVPD.

<table>
<tr><th rowspan="2">System</th><th rowspan="2">$\Delta G_{prot}$</th><th rowspan="2">$TS_{add}$ (1)</th><th colspan="2">$TS_{elim}$ (2)</th><th rowspan="2">$Int^{+/-}$</th></tr>
<tr><th>$TS_{PT}$(2)</th><th>$TS_{elim}$(3)</th></tr>
<tr><td rowspan="2">Lys·CAM2($ImH^+$)·$H_2O$</td><td rowspan="2">+8.8</td><td rowspan="2">15.1</td><td colspan="2">15.3</td><td rowspan="2">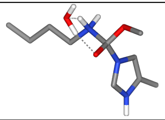</td></tr>
<tr><td>21.7</td><td>14.2</td></tr>
<tr><td rowspan="2">Lys·CAM2($ImH^+$)·$2H_2O$</td><td rowspan="2">+8.8</td><td rowspan="2">13.5</td><td colspan="2">13.5</td><td rowspan="2">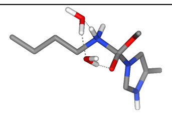</td></tr>
<tr><td>17.1</td><td>15.1</td></tr>
<tr><td rowspan="2">Lys·CAM2($ImH^+$)·Arg·$H_2O$</td><td rowspan="2">+8.8</td><td rowspan="2">11.3</td><td colspan="2">11.9</td><td rowspan="2">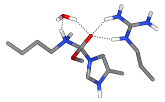</td></tr>
<tr><td>21.5</td><td>14.4</td></tr>
<tr><td rowspan="2">Lys·CAM2($Im^0$)·Arg·$H_2O$</td><td rowspan="2">+5.2</td><td rowspan="2">12.5</td><td colspan="2">-</td><td rowspan="2">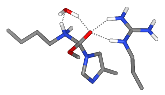</td></tr>
<tr><td>23.0</td><td>-</td></tr>
<tr><td rowspan="2">Lys·CAM2($ImH^+$)·Arg·$H_2O$</td><td rowspan="2">+8.8</td><td rowspan="2">12.4</td><td colspan="2">11.6</td><td rowspan="2">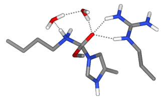</td></tr>
<tr><td>14.2</td><td>17.3</td></tr>
<tr><td>Lys·CAM2($ImH^+$)·Asp</td><td>+8.8</td><td>13.3</td><td colspan="2">7.8*</td><td>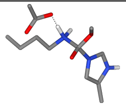</td></tr>
<tr><td>Lys·CAM2($ImH^+$)·Asp·$H_2O$</td><td>+8.8</td><td>11.9</td><td colspan="2">6.8*</td><td>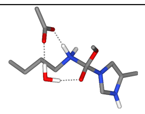</td></tr>
<tr><td>Lys·CAM2($ImH^+$)·Arg·Asp</td><td>+8.8</td><td>12.7</td><td colspan="2">13.1</td><td>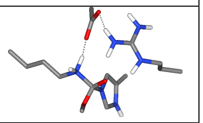</td></tr>
</table>

### List of Figures


S1 Energy profile of Lys labeling with both Im ring protonation states. $TS_{elim}$* indicates the energy barrier of the precursor step that directly triggers the subsequent elimination. . . . . . . . . . . . . . . . . . . . . . . . . . . . . S-2

S2 Linear regression for the determination of the $pK_a$ of the cropped-CAM2 Im ring. Geometry and frequency calculations were done with the B3LYP/def2-SVPD, employing the optimized geometries to perform single-point energy evaluations at DSD-PBEP86/def2-TZVPD. . . . . . . . . . . . . . . . . . . . . S-3

S3 Energy profile of CAM2 hydrolysis with both Im ring protonation states. . . S-4

S4 Fractions of hydrogen bond between PFQX antagonist and GluA2(S1S2J) protein for each monomer and replicate. Diagonal hatch (// and \\) distinguishes the monomers. . . . . . . . . . . . . . . . . . . . . . . . . . . . . . . S-5

S5 Time series of PFQX antagonist binding measured as the distance between GluA2(S1S2J) Arg96(C$\zeta$) and the center of mass of the quinoxalinedione oxygens for all five replicates and each monomer. . . . . . . . . . . . . . . . . S-6

S6 Time series of the top (upstream) and bottom (downstream) distances with respect the membrane in the whole protein. Top distance as the distances between monomer-monomer Ser229(C$\alpha$) and bottom distance as the distances between Lys185(C$\alpha$). Gray horizontal lines indicate the initial values of both distances. . . . . . . . . . . . . . . . . . . . . . . . . . . . . . . . . . . S-7

S7 Time series of clamshell state measured as the distance between the center of mass of Leu90, Thr91 and Ile92 (S1 domain) and Ser142 and Thr143 (S2 domain). . . . . . . . . . . . . . . . . . . . . . . . . . . . . . . . . . . . S-8

S8 Root mean square deviation between monomer-monomer main chains for all five replicates. . . . . . . . . . . . . . . . . . . . . . . . . . . . . . . . . . S-9

S9 Time series and density distributions of the distance between the Lys60 and Ser140 side chains for each monomer in the five replicates. The black and yellow dashed line indicates the distance in the case of a one-water bridge. . S-10

S10 Time series and density distributions of the distance between the Lys60 and Asp58 side chains for each monomer in the five replicates. The black horizontal line indicates the distance in the case of a hydrogen bond. . . . . . . . . . . S-11

S11 Time series and density distributions of the distance between the Ser14 and Glu13 side chains for each monomer in the five replicates. The black horizontal line indicates the distance in the case of a hydrogen bond. . . . . . . . . . . S-12

S12 Time series and density distributions of the distance between the Ser140 and Asp139 side chains for each monomer in the five replicates. The black and yellow dashed line indicates the distance in the case of a hydrogen bond. . . S-13

S13 Time series of antagonist-moiety binding measured as the distance between Arg96(C$\zeta$) and the center of mass of the CAM2 quinoxalinedione oxygens for all nine replicates and each monomer of C1 set. . . . . . . . . . . . . . . . . . S-14

S14 Time series of antagonist-moiety binding measured as the distance between GluA2(S1S2J) Arg96(C$\zeta$) and the center of mass of the CAM2 quinoxalinedione oxygens for all nine replicates and each monomer of C2 set. . . . . . . S-15

S15 Time series of the top (upstream) and bottom (downstream) distances with respect the membrane in the whole protein. Top distance as the distances between monomer-monomer Ser229(C$\alpha$) and bottom distance as the distances between Lys185(C$\alpha$) of C1 set. Black horizontal lines show the initial distances for both sets; the current set is represented by the solid line. . . . . S-16

S16 Time series of the top (upstream) and bottom (downstream) distances with respect the membrane in the whole protein. Top distance as the distances between monomer-monomer Ser229(C$\alpha$) and bottom distance as the distances between Lys185(C$\alpha$) of C2 set. Black horizontal lines show the initial distances for both sets; the current set is represented by the solid line. . . . . . S-17
S17 Top and bottom distances of C1 and C2 GluA2(S1S2J)-CAM2 sets onto the 2D clustering projection from GluA2(S1S2J)-PFQX simulations. . . . . . . . S-18
S18 1D distributions of the distances between each labeled nucleophile and the acyl carbon electrophile for both C1 and C2 sets. . . . . . . . . . . . . . . . . S-18
S19 2D distances distributions between CAM2 acyl cabron and Lys60(N$\zeta$) and Ser140(O$\gamma$) for all nine replicates of C1(above) and C2(below) sets. Same data is plotted from left to right but employing the color to indicate the respective distance with Ser14(O$\gamma$), the replicates and monomers populations and, the data density. × represents a high-density population present in set C2 but not in C1, identified as a geometry corresponding to a possible critical point where Ser140 is positioned to facilitate the labeling of Lys60. . . . . . S-19
S20 Barplots of fractions of hydrogen bond interactions of CAM2 with GluA2(S1S2J) protein. Graph is separated in three parts of CAM2, from above to below: the antagonist fragment (QX), the middle that include amide groups (am1-3) and reactive acyl and, Im groups and the oregon green (OG) fragment. C2 is distinguished from C1 by diagonal hatching (//). . . . . . . . . . . . . . . . . S-20
S21 Barplots of fractions and presences of hydrogen bonds between GluA2(S1S2J) and CAM2 when each nucleophile Lys(N$\zeta$), Ser14(O$\gamma$) and Ser140(O$\gamma$) are at least 5 Å from the CAM2 acyl carbon. Hydrogen bond interactions are ordered according CAM2 atoms from antagonist fragment (QX) to Oregon Green (OG). CAM2 amide groups are indicate as am1-3. Side hydrogen bond interactions involving residues that interact with CAM2 are also shown. . . . S-21
